# Elucidating Molecular Mechanisms of Protoxin-2 State-specific Binding to the Human Na_V_1.7 Channel

**DOI:** 10.1101/2023.02.27.530360

**Authors:** Khoa Ngo, Diego Lopez Mateos, Yanxiao Han, Kyle C. Rouen, Surl-Hee Ahn, Heike Wulff, Colleen E. Clancy, Vladimir Yarov-Yarovoy, Igor Vorobyov

## Abstract

Human voltage-gated sodium (hNa_V_) channels are responsible for initiating and propagating action potentials in excitable cells and mutations have been associated with numerous cardiac and neurological disorders. hNa_V_1.7 channels are expressed in peripheral neurons and are promising targets for pain therapy. The tarantula venom peptide protoxin-2 (PTx2) has high selectivity for hNa_V_1.7 and serves as a valuable scaffold to design novel therapeutics to treat pain. Here, we used computational modeling to study the molecular mechanisms of the state-dependent binding of PTx2 to hNa_V_1.7 voltage-sensing domains (VSDs). Using Rosetta structural modeling methods, we constructed atomistic models of the hNa_V_1.7 VSD II and IV in the activated and deactivated states with docked PTx2. We then performed microsecond-long all-atom molecular dynamics (MD) simulations of the systems in hydrated lipid bilayers. Our simulations revealed that PTx2 binds most favorably to the deactivated VSD II and activated VSD IV. These state-specific interactions are mediated primarily by PTx2’s residues R22, K26, K27, K28, and W30 with VSD as well as the surrounding membrane lipids. Our work revealed important protein-protein and protein-lipid contacts that contribute to high-affinity state-dependent toxin interaction with the channel. The workflow presented will prove useful for designing novel peptides with improved selectivity and potency for more effective and safe treatment of pain.

**Summary:** Na_V_1.7, a voltage-gated sodium channel, plays a crucial role in pain perception and is specifically targeted by PTx2, which serves as a template for designing pain therapeutics. In this study, *Ngo et al.* employed computational modeling to evaluate the state-dependent binding of PTx2 to Na_V_1.7.

## 1. Introduction

Voltage-gated sodium (Na_V_) channels are responsible for electric signaling and play a key role in various physiological processes^1,2^. Na_V_ channels are involved in the functioning of the central and peripheral nervous systems and the contraction of skeletal and cardiac muscles^2^. There are nine known members in the Na_V_ family, named Na_V_1.1 to Na_V_1.9^2^. A pore-forming α subunit or a eukaryotic Na_V_ channel is composed of four homologous domains (DI-DIV), each containing six transmembrane segments (S1-S6), with S1 through S4 serving as the voltage-sensing domain (VSD). The S4 segment contains four or five positively charged amino acid residues known as gating charge residues and acting together as a voltage sensor. The S5 and S6 segments, along with intervening loops and pore helix form the channel pore which contains the ion selectivity filter (SF) region. Membrane depolarization induces the charged residues in the S4 segment to move outwards, a movement that is transmitted to the pore domain through the S4-S5 linker leading to channel opening and a rapid influx of sodium ions into the cell. This sodium influx further depolarizes the cell membrane and leads to the generation of an action potential. The opening of Na_V_ channels is dependent on the sequential activation of VSD I through III^3,4^. In contrast, the activation of VSD IV is coupled with the rapid inactivation of the channel through the release of the isoleucine – phenylalaine – methionine (IFM) or similar motif in the DIII-DIV intracellular loop (inactivation gate)^3–6^. This inactivation process is crucial to stop the influx of sodium ions and to enable the repolarization of the membrane to terminate the action potential.

Na_V_ channel dysfunction arising from mutations can lead to various pathophysiological conditions including arrhythmias, epilepsy, chronic pain, and insensitivity to pain^7^. Na_V_1.3, Na_V_1.7, Na_V_1.8, and Na_V_1.9 channels have been implicated in pain signaling^2^. Transgenic mice lacking Na_V_1.7- and Na_V_1.8-positive nociceptors have been shown in experiments to exhibit increased mechanical and thermal pain thresholds^8^. In particular, the mice showed reduced or no response to inflammatory pain responses evoked by various stimuli^8^. Moreover, humans born with loss-of-function mutations in the SCN9A gene encoding for the Na_V_1.7 α subunit were discovered to have congenital insensitivity to pain^9^. In contrast, gain-of-function mutations were associated with aberrant chronic pain sensation and related disorders such as inherited erythromelalgia and paroxysmal extreme pain disorder^10,11^. These observations further emphasize the importance of the Na_V_1.7 channels to pain perception.

Local anesthetics, such as lidocaine, are used to reduce the transmission of pain signals to the central nervous system by non-selectively blocking various Na_V_ subtypes^12^. Due to their lack of Na_V_ selectivity, side effects are often reported concerning the loss of other sensations, such as the sense of touch. Use of opioid analgesics is associated with side effects such as respiratory depression, constipation, and development of dependence^13^. Therefore, developing drugs that selectively inhibit Na_V_1.7 function may result in strong analgesia without undesirable side effects^14,15^. Na_V_ channels implicated in pain signaling have been discovered to be targets of natural toxins found in species such as spiders and scorpions^7^. The toxins modulate channel activity by either blocking the ion permeation pathway (i.e., pore blockers) or binding to the VSD to alter channel gating kinetics (i.e., gating modifier toxins)^16^.

Protoxin-2 (PTx2), a 30-residue peptide derived from the venom of the Peruvian Green Velvet tarantula (*Thrixopelma pruriens*), is a gating modifier toxin and has a moderate selectivity for Na_V_1.7 versus other Na_V_ subtypes, making it a suitable scaffold for a peptide-based pain therapeutic^17^. This toxin interferes with the activation of Na_V_ channels by binding to the S3-S4 loop in the deactivated VSD II and shifting the voltage dependence of activation to a more positive potential^17^. PTx2 is 100- fold more potent for Na_V_1.7 (IC_50_ = 0.3-1.0 nM) versus other Na_V_ subtypes^14,18–20^. Additionally, PTx2 binds to the deactivated hNa_V_1.7 VSD IV inhibiting fast inactivation and producing sustained sodium currents, similar to the effect of α-scorpion toxins^4^, but with lower potency (IC_50_ = 0.24 μM)^19^. However, this effect of PTx2 is typically masked by the preferential inhibition of Na_V_ channel activation due to higher affinity binding to VSD II^19^.

By using peptide toxins as a scaffold for designing novel ion channel modulators, researchers can potentially develop more effective and targeted therapies for chronic pain^15,21^. Potential strategies for pain therapy via hNa_V_1.7 gating inhibition include the trapping of VSD II in the deactivated state^22^ (to prevent the channel from activating) or of VSD IV in the activated state^23^ (to keep the channel in the inactivated, non-conducting state). However, current structural studies^4,17^ have not yet resulted in a complete atomic-resolution understanding of how PTx2 interacts with Na_V_1.7 VSDs in a state-specific manner as well as with the surrounding lipids. Recently, experimental structures of PTx2 bound to Na_V_1.7 VSD II in various states have been resolved using chimeric constructs, grafting parts of the human Na_V_1.7 channel sequence into a bacterial Na_V_ channel^22^. These studies resulted in X-ray and cryo-EM structures of the PTx2-hNa_V_1.7-VSD II-Na_V_Ab complex in activated and deactivated states^22^. Yet, these structures do not fully represent a human Na_V_ channel. Notably, Shen *et al.*^17^ obtained the first cryo-EM structure of the full hNa_V_1.7, resolving structures of PTx2 bound to activated VSD II and VSD IV. However, due to the relatively low resolution of PTx2 densities, these structures did not allow complete atomic reconstruction of the peptide toxin. In addition, the studies did not provide energetic insights into toxin-channel/lipids interactions, which are crucial to designing improved peptides as ion channel modulators.

Computational methods can provide structural insights into the atomic-level mechanisms involved in ion channel function and modulation^24^ and toxin binding^25–27^. Moreover, they are useful tools to guide the rational design process towards enhanced peptide variants^14,28^. In our study, we demonstrated that computational structural modeling and molecular dynamics simulations could accurately capture the atomic-resolution molecular mechanisms by which PTx2 binds to hNa_V_1.7 VSD II and IV in the activated and deactivated states. Our molecular modeling and simulation insights will be useful to inform future peptide design studies and help researchers to identify potential peptide mutations to improve PTx2 selectivity and potency for hNa_V_1.7. Moreover, our methodology can be expanded to study other toxin-channel interactions to design potential peptide-based therapeutics for diseases such as cancer^29^, cardiac arrhythmia^30^, epilepsy^30^, and neuropathic pain^30^.

## 2. Results

### 2.1 General overview of different PTx2 - Na_V_1.7 VSD structures

In this study, we employed computational modeling, informed by experimental structural data, to investigate the molecular interactions between PTx2 and hNa_V_1.7’s Voltage Sensor Domains (VSDs) II and IV in various states (Figure 1). It should be noted that our focus was solely on individual VSDs and not the whole channel alpha subunit, an approach selected to optimize the use of computational resources. While experimental structures of activated/deactivated VSD IV and activated VSD II of human Na_V_1.7 are available^17,22^, the experimental structure of deactivated VSD II of hNa_V_1.7 has yet to be determined. To acquire the necessary structural information, we sought to obtain structures of both hNa_V_1.7 VSD II and VSD IV in both activated and deactivated conformations. To obtain a structure of deactivated VSD II, we used the experimental structure of the hNa_V_1.7-VSD II-Na_V_Ab^22^ as a template for homology modeling using the RosettaCM^31^ (sequence identity: 63.7%). We then compared the positions of gating charges in each VSD by calculating the distances between equivalent C_α_ atoms after VSD superimposition (**Figure 10**). Our analysis revealed that gating charges of VSD II move ∼8.5 Å, on average, between the deactivated and activated states. In the deactivated state, only the first gating charge (“R1”) is above the hydrophobic constriction site (HCS), while in the activated state “R1”, “R2” and “R3” are above the HCS. VSD IV showed a more dramatic displacement of the gating charges (∼13.8 Å) between the activated and deactivated states. Just as in VSD II, in the deactivated conformation, only “R1” is located above the HCS; however, in the activated conformation of VSD IV “R4” also reaches above the HCS, which contrasts with the activated VSD II structure where only “R1” to “R3” lie above the HCS. These results suggest that the VSD II in the Na_V_1.7 cryo-EM structure (PDB: 6J8J^17^) is not fully activated, which might be explained by the presence of PTx2 or the lack of membrane voltage during the structure determination process.

**Figure 1.**
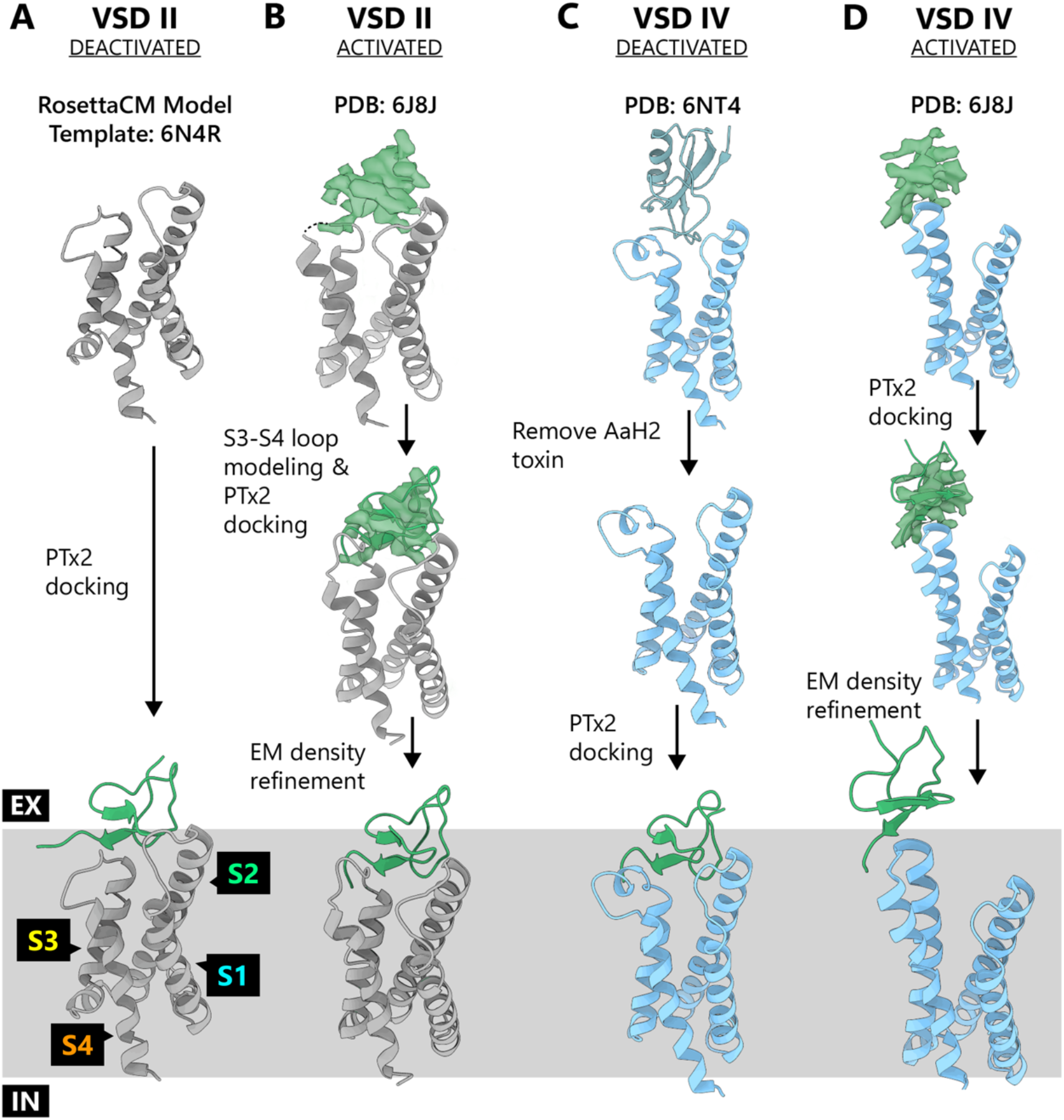
Workflow of the structural refinement, modeling, and docking process for different PTx2 – hNa_V_1.7 VSD II and IV structures. The membrane is represented as a gray box, with the extracellular- and intracellular-facing sides labeled as "EX" and "IN" respectively.

Subsequently, structural models of PTx2 in complex with VSD II and VSD IV were generated in both deactivated and activated states as starting points in MD simulations. The methodology utilized for the generation of these models is outlined in **Figure 1** and briefly summarized as follows: RosettaDock with a membrane energy function was employed for docking PTx2 to the VSDs. The activated conformations of the VSDs (PDB: 6J8J^17^) with bound PTx2 were further refined using the available experimental EM density maps.

Previously, it was determined that PTx2 interferes with Na_V_1.7 activation by binding potently (IC_50_ = 0.3-1.0 nM) to the S3-S4 loop in VSD II and shifting the voltage dependence of activation to more depolarized potentials^14,17–20^. Analysis of the structural models generated here provided important insights into the conformational-dependent binding of PTx2 to Na_V_1.7 VSD II & VSD IV. PTx2 binds to the VSD II at the “LFLAD” motif, which comprises residues L812 to D816 and is located within the S3 segment and S3-S4 loop^22^. The toxin binding interface can be divided into two distinct regions: polar and hydrophobic. In the PTx2 - deactivated VSD II model (**Figure 2A, panel i**), the first gating charge (“R1”) is observed to be located below the channel-toxin polar binding interface. The electronegative pocket created by VSD II residues E811 and D816 forms hydrogen bonds and salt bridges with the positively charged toxin residues K26 and R22 (**Figure 2A, panel iii**). The hydrophobic interface (**Figure 2A, panel ii**) is composed of M6 and tryptophan residues W5, W24, and W30 from the toxin and L812 and F813 from the channel’s VSD II S3 segment. Additionally, W24 was observed to be positioned into the VSD top wedge, making hydrophobic contacts with channel residues A776 and L770 from VSD II segment S2 (**Figure 2A, panel iv**).

**Figure 2.**
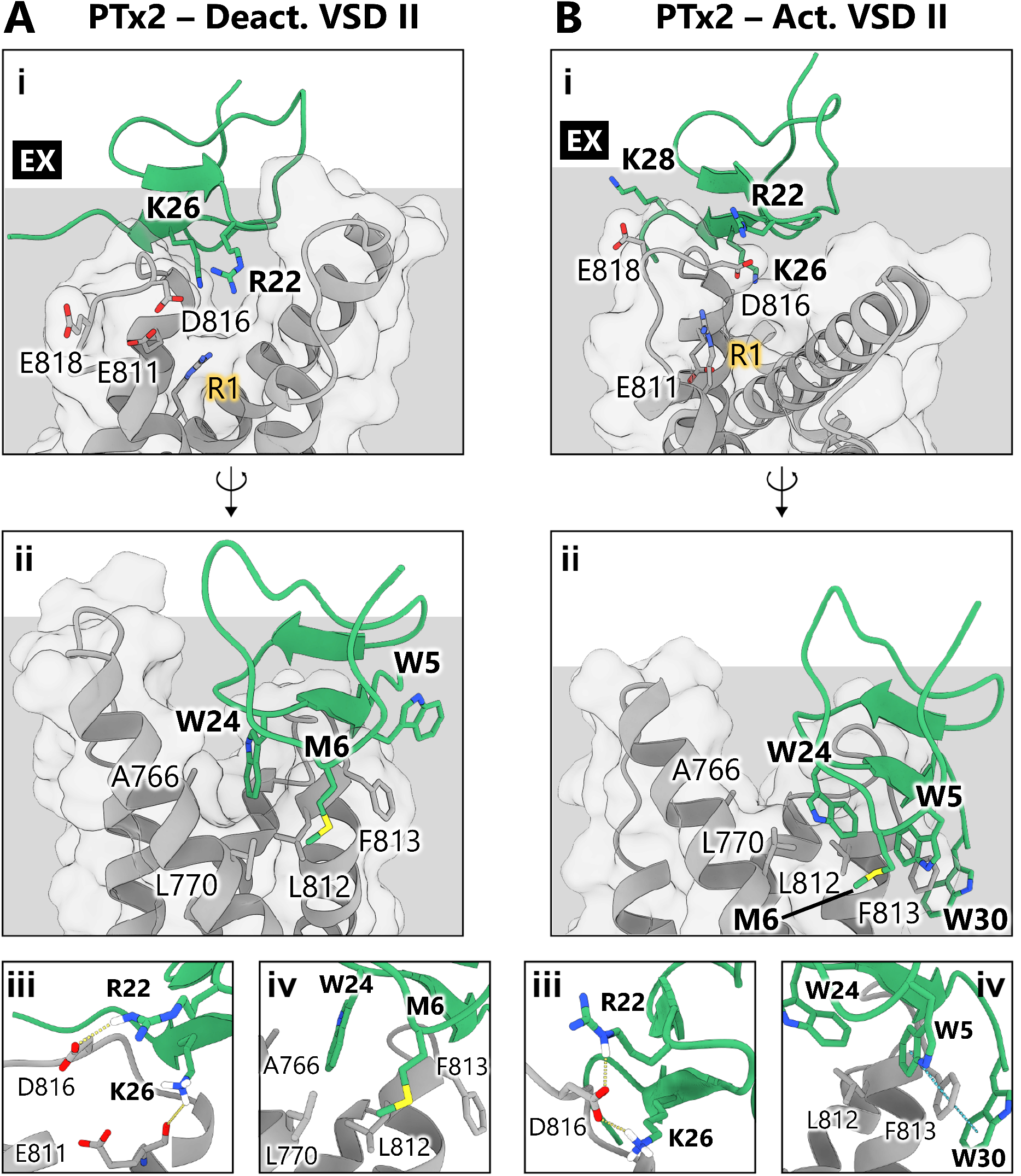
Structural models of PTx2 bound to hNa_V_1.7 VSD II in the deactivated and activated states. Binding interface of PTx2 – deactivated VSD II (A) and PTx2 – activated VSD II (B) complexes of top scoring Rosetta models. i) Front view of interface the complex highlighting interactions with the VSD that comprise the polar binding interface; “R1” highlights the position of the first gating charge. ii) Back view of the complex highlighting interactions with the VSD that comprise the hydrophobic or membrane-interacting interface. iii) detailed interactions at the polar interface; yellow dashed lines highlight hydrogen bonds. iv) detailed interactions at the hydrophobic interface; blue dashed lines highlight π - stacking interactions. The membrane is represented as a gray box, with the extracellular-facing side labeled as "EX".

In the PTx2 – activated VSD II model (**Figure 2B**), the upward movement of the S4 segment and subsequent relocation of the “R1” gating charge residue disturbs the electronegative pocket, and the toxin experiences a global shift that pulls it away from the VSD (**Supplementary Video 1**). The main interaction in our PTx2 – activated VSD II model at the polar interface is between toxin R22 and K26 residues and channel D816 residue (**Figure 2B, panel iii**). Also, channel’s residue E818 interacts with K28 from the toxin, although this interaction is not present in any of the experimental structures. The hydrophobic interface (**Figure 2B, panel ii**) comprises π-stacking interactions between toxin W5 and W30 residues and channel F813 residue (**Figure 2B, panel iv**). The overall shift of the toxin resulted in toxin’s residue W24 no longer interacting with channel’s residues of VSD II S2 segment.

Experimental results show that PTx2 can also bind to Na_V_1.7 VSD IV and modulate channel inactivation^19^, although with much lower potency (IC_50_ = 0.24 μM). Our models of PTx2 bound to both deactivated and activated VSD IV presented binding interfaces, which can also be separated into a polar interface and a hydrophobic interface. Our models revealed that the position and interface of PTx2 changed significantly more between the deactivated and activated states of VSD IV compared to VSD II (**Supplementary Video 2**).

Our model of PTx2 bound to the deactivated VSD IV was built using the experimental structure of the hNa_V_1.7 VSD IV in a deactivated state induced by binding of an α-scorpion toxin (PDB: 6NT4^4^), and it is important to note that the actual VSD IV deactivated state to which PTx2 binds may differ. In our final top Rosetta model, which was selected for subsequent simulations, PTx2 is embedded in the membrane between the VSD IV S3 and S2 segments. hNa_V_1.7 VSD IV residues D1586 and E1589 and K26 from the toxin (**Figure 3A, panel i**). Toxin R22 extends into the VSD and establishes hydrogen bonds with channel’s residue D1586 while K26 interacts with G1581 at the top of the S3 segment (**Figure 3A, panel iii**). The hydrophobic interface is more complex in this case (**Figure 3A, panel ii**). PTx2 makes contacts with residues in the VSD IV S2 segment, which are mainly hydrophobic interactions mediated by toxin residue W24 and channel residues Y1539 and W1538. Finally, toxin residues W5, M9, and W30, which also serve as membrane anchors, interact with several hydrophobic residues from the channel located in the VSD IV S3 segment: M1582, F1583 and L1584 (**Figure 3A, panel iv**). In our models, a large conformational rearrangement of the VSD IV occurs between the deactivated and the activated state. As a result, the location of PTx2 also undergoes a dramatic change (**Supplementary Video 2**).

**Figure 3.**
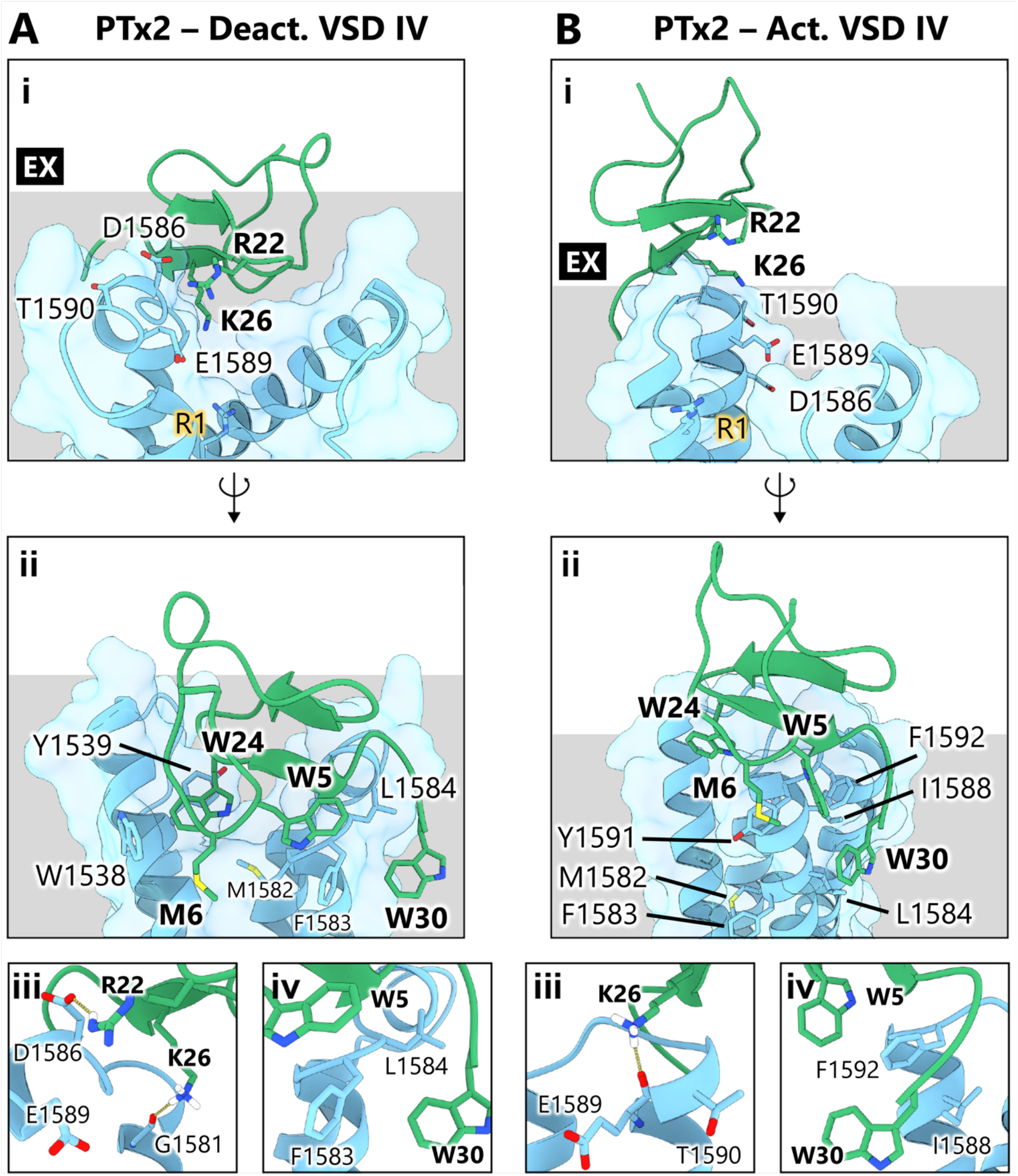
Structural models of PTx2 bound to hNa_V_1.7 VSD IV in the deactivated and activated states. Binding interface of PTx2 – deactivated VSD IV (A) and PTx2 – activated VSD IV (B) complexes of top-scoring Rosetta models. i) Front view of the complex highlighting interactions with the VSD that comprise the polar binding interface; “R1” highlights the position of the first gating charge. ii) Back view of the complex highlighting interactions with the VSD that comprise the hydrophobic or membrane-interacting interface. iii) Detailed interactions at the polar interface; yellow dashed lines highlight hydrogen bonds. iv) Detailed interactions at the hydrophobic interface. The membrane is represented as a gray box, with the extracellular-facing side labeled as "EX".

In the activated VSD IV state, the toxin is situated at the top of the S3 segment and does not establish interactions with the S2 segment. The polar interface observed in the activated VSD IV state (**Figure 3B, panel i**) is primarily mediated by the interaction between toxin residue K26 and channel’s S3 residue E1589 (**Figure 3B, panel iii**), with PTx2 residue R22 not appearing to play a significant role in this interaction in contrast to the deactivated state. In the hydrophobic interface (**Figure 3B, panel ii**), the toxin uses its residues W5, M9, and W30 to anchor to the membrane and establish hydrophobic contacts with residues I1588, Y1591, F1592 from the channel’s S3 segment (**Figure 3B, panel iv**). Notably, these hydrophobic channel VSD IV residues are positioned far from the binding interface in the deactivated state. In contrast, the channel residues previously involved in the hydrophobic interface in the deactivated state (M1582, F1583 and L1584) now lie below the interface and do not interact with the toxin (**Figure 3B, panel ii**).

### 2.2 Molecular dynamics simulations on PTx2 interactions with hNa_V_1.7 VSD II and IV

We conducted molecular dynamics (MD) simulations on the docked structures to assess their stability in an environment more closely resembling physiological conditions. These simulations, carried out in three independent 1-µs long replicates, were instrumental in quantifying the energetics of the binding process and investigating the impact of lipids on binding. The simulations showed converging behavior during these timeframes, the details of which are discussed further below. To analyze the simulations, we utilized the atomic coordinates from the entire simulation trajectory, excluding 90-ns long equilibration phase. From this data, contact maps were generated, revealing the residues involved in PTx2 – hNa_V_1.7 VSDs binding interactions, along with the type (e.g., hydrophobic, hydrogen bond, salt bridges, π-stacking, or cation-π) and duration of these interactions. The criteria used to quantify these interactions are indicated in **Supplementary Table S1**. The contact maps are displayed in **Supplementary Figures S1-S4**.

**PTx2**–**hNa_V_1.7 deactivated and activated VSD II:** Throughout the simulation, PTx2 consistently remained bound to VSD II by lodging itself in a cleft located between the S1-S2 and S3-S4 loops, as depicted in **Figure 4A, D**. Hydrophobic contacts constitute the majority of interactions between the toxin and deactivated or activated VSD II (57 ± 4% or 55 ± 5%), followed by hydrogen bond formation (30 ± 2% or 29 ± 6%) and salt bridges (13 ± 2% or 16 ± 8%), as shown in **Figure 4B, E**.

**Figure 4.**
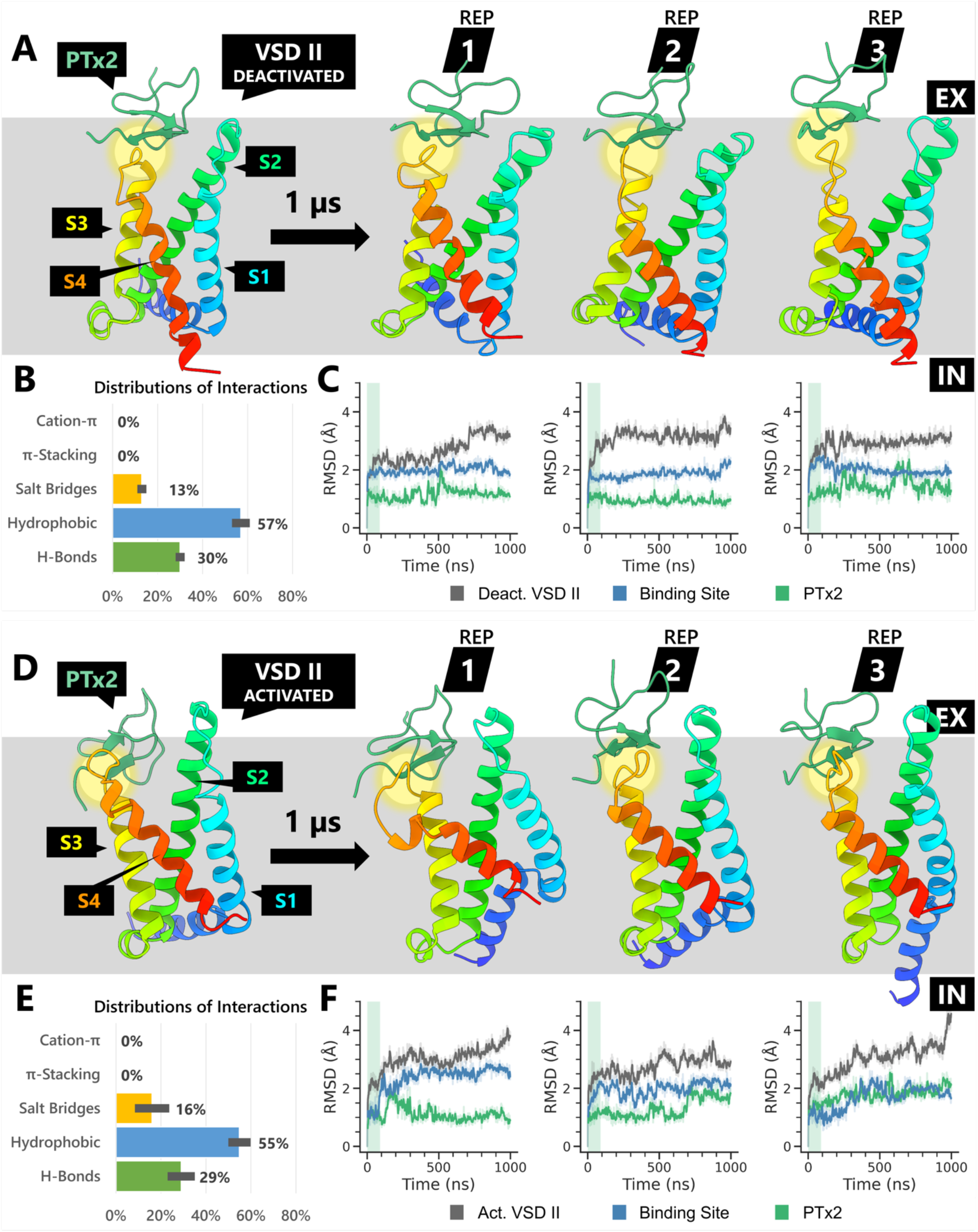
General overview of the three replicas (REP) of 1-μs molecular dynamics simulations of PTx2 – hNa_V_1.7 VSD II in the deactivated (A, B, C) and activated states (D, E, F). The lipid membrane is represented as a gray box, with the extracellular- and intracellular-facing sides labeled as "EX" and "IN" respectively. PTx2 – VSD binding sites are highlighted in yellow circles. A, D) Conformational changes of the toxin – VSD complex following the 1-μs long molecular dynamics simulation for each replica. B, E) Distribution of all toxin – VSD interaction types encountered during the simulations, computed by dividing the sum of a specific interaction type by the sum of all interactions across all simulation frames. C, F) Time series of the root-mean-square deviation (RMSD) of PTx2, entire VSD and its toxin binding site backbone atoms compared to the initial structures. The simulation equilibration period is highlighted in green.

The backbone RMSD profiles, as depicted in **Figure 4C** and **6F**, suggest that the toxin underwent only minor structural changes during the 1-μs simulations, approximately between 1 to 2 Å RMSD. In contrast, the VSDs underwent more considerable structural alterations during the simulations, about 3 to 4 Å RMSD. The instability could be due to their disconnection from the rest of the channel in addition to the flexibility of the loops as well as the pre-S1 helix. However, the RMSD of the toxin-binding region, defined as the area within a 7.5 Å radius of where the toxin interacts with the VSD, experienced only minor fluctuations, approximately between 1 to 2.5 Å RMSD.

**Supplementary Figures S1 and S2** present the contact maps, averaged from three replicas, showcasing the interactions between PTx2 and VSD II in the deactivated and activated states, respectively, along with any interactions with lipids. In both cases, PTx2 formed hydrogen bonds, hydrophobic interactions, and salt bridges primarily with residues on the S3-S4 loop of VSD II (including E811, L812, F813, L814, A815, D816), as depicted in **Figure 5**.

**Figure 5.**
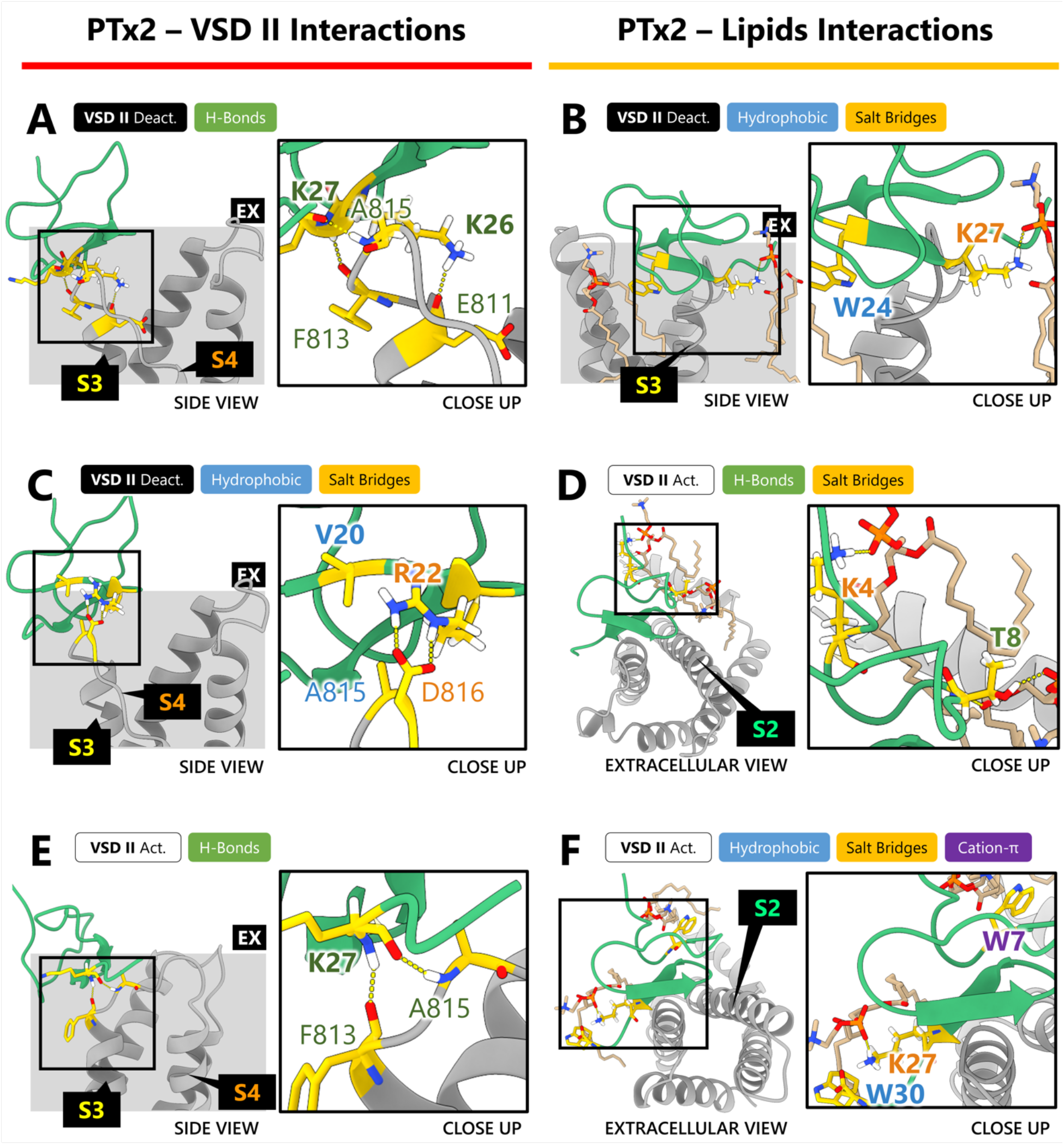
Dominant interactions recorded from molecular dynamics simulations between PTx2 and hNa_V_1.7 VSD II (A, C, E) or PTx2 and POPC membrane (B, D, F). Residues engaged in binding are highlighted in gold. PTx2 is represented in green, while the VSD is depicted in gray. Each panel features interaction types presented in color-coded boxes whose color corresponds to the involved residues’ labels. The structures were visualized at the end of the 1-μs long MD simulations.

In most cases, the root-mean-square deviation (RMSD) of the PTx2 binding site showed stabilization during the latter half of the simulation (≥ 500 ns). Based on this observation, we classified a contact between two residues as dominant if the average contact duration derived from three replicas, plus standard errors of the mean, exceeds 75% (or 50% for protein-lipid interactions) of the time in this latter half of the simulation. Our subsequent analyses primarily centered on characterizing and visualizing these dominant interactions that drove the binding process. That said, we also paid attention to any noteworthy non-dominant contacts that emerge in the study.

During PTx2 binding to the deactivated VSD II, several key interactions were observed. PTx2 K26 formed a hydrogen bond with VSD II E811 carbonyl located on the S3 loop, while K27 backbone engaged in hydrogen bonding with F813 carbonyl and A815 amide on the VSD II S3-S4 loop (**Figure 5A**). PTx2 V20 participated in a hydrophobic interaction with VSD II residue A815, and toxin’s residue R22 formed a salt bridge with VSD II residue D816, as depicted in **Figure 5C**. Notably, as shown in **Figure 5B**, PTx2 was observed to anchor itself to the membrane through hydrophobic interactions involving residue W24 and the surrounding lipids. Additionally, its residue K27 formed salt bridges with nearby lipid head groups. In addition to these dominant interactions, **Supplementary Figure 1** shows toxin’s residues L23 and W24 also sporadically engaged in hydrophobic interactions with VSD II S1-S2 residues N763, I767, and L770, effectively creating two anchors that positioned the toxin between the VSD loops.

Intriguingly, when PTx2 binds to the activated VSD II, it exhibited a stronger reliance on interactions with the surrounding lipids rather than VSD II residues. Similar to the previous case, toxin’s residue K27 backbone engaged in hydrogen bonding with F813 carbonyl and A815 amide on the S3-S4 loop (**Figure 5E**). As shown in **Figure 5D**, the PTx2 T8 side chain hydroxyl group engaged in hydrogen bonding and to a lesser extent, the K4 side chain amino group formed a salt bridge with nearby POPC phosphate (PO4) groups. Moreover, as shown in **Figure 5F**, toxin’s residue W30 engaged in hydrophobic interactions with lipids and to a lesser extent, K27 created a salt bridge with POPC PO4 groups near the VSD II S3 location. At the same time, PTx2 residue W7 participated in cation-π interactions with neighboring lipids close to VSD II S2 segment. Such interactions contributed to the formation of hydrophobic anchors that helped orient the toxin binding to the VSD II. Although to a lesser extent than the previous case, some hydrophobic interactions were observed involving PTx2 residues W5 and V2 with VSD II residues F813 and A815 (**Supplementary Figure 2**). Additionally, toxin’s R22 sporadically formed salt bridges with VSD II residues E753 (S1-S2 loop) and D814 (S3-S4 loop).

**PTx2**–**hNa_V_1.7 deactivated and activated VSD IV:** PTx2 exhibits a dynamic range of binding poses when interacting with VSD IV in the deactivated and activated states. During the simulation, PTx2 progressively adjusted its binding pose, positioning itself deeper into the binding pocket nestled between the S1-S2 and S3-S4 loops as displayed in **Figure 6A**. Hydrophobic contacts formed most of the interactions between PTx2 and the deactivated VSD IV, accounting for 45 ± 5% of the total as demonstrated in **Figure 6B**. These are followed by hydrogen bonds and salt bridges, constituting 32 ± 1% and 22 ± 5% of the interactions respectively. Throughout the simulations, **Figure 6C** depicts considerable fluctuations in the backbone RMSD profiles of the VSD and the binding site, varying between 2 to 3.5 Å. This significant variability primarily resulted from the flexible S3-S4 loop’s movement as PTx2 embedded itself into the crevice formed between the two VSD loops.

**Figure 6.**
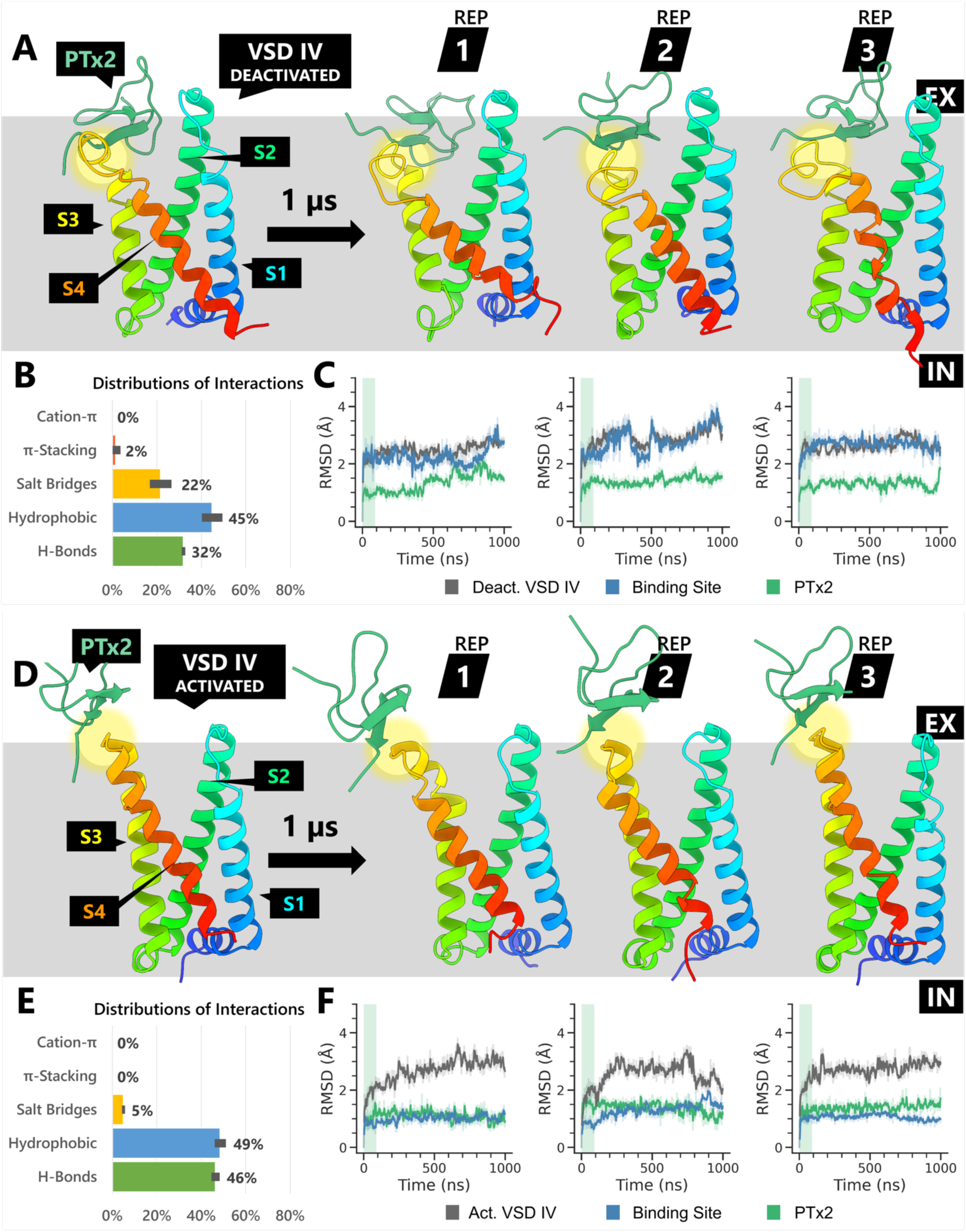
A general overview of the three replicas (REP) of 1-μs molecular dynamics simulations of PTx2 – hNa_V_1.7 VSD IV in the deactivated (A, B, C) and activated states (D, E, F). The lipid membrane is represented as a gray box, with the extracellular- and intracellular-facing sides labeled as "EX" and "IN" respectively. PTx2 – VSD binding sites are highlighted in yellow circles. A, D) Conformational changes of the toxin – VSD complex following the 1-μs long molecular dynamics simulation for each replica. B, E) Distribution of all toxin – VSD interaction types encountered during the simulations, computed by dividing the sum of a specific interaction type by the sum of all interactions across all simulation frames. C, F) Time series of the root-mean-square deviation (RMSD) of PTx2, entire VSD and its toxin binding site backbone atoms compared to the initial structures. The simulation equilibration period is highlighted in green.

**Supplementary Figures S3 and S4** present the contact maps, averaged from three replicas, showcasing the interactions between PTx2 and VSD IV in the deactivated and activated states, respectively, along with any interactions with lipids. Throughout the simulation, PTx2 was bound to the deactivated VSD IV via a multitude of interactions with the residues on the S3-S4 loop of the VSD. As depicted in **Figure 7A**, PTx2’s K27 formed several hydrogen bonds with F1583 carbonyl and A1585 amide backbone groups, both situated at the S3 helix’s terminus of VSD IV where the S3-S4 loop commences. Toxin’s residues K26 and K28 established salt bridges with VSD IV residues E1589 and D1586, respectively, located higher on the S3-S4 loop, as indicated in **Figure 7B**. Additionally, PTx2’s R22 contributed to salt bridge formation with D1586, albeit to a lesser extent than K28.

**Figure 7.**
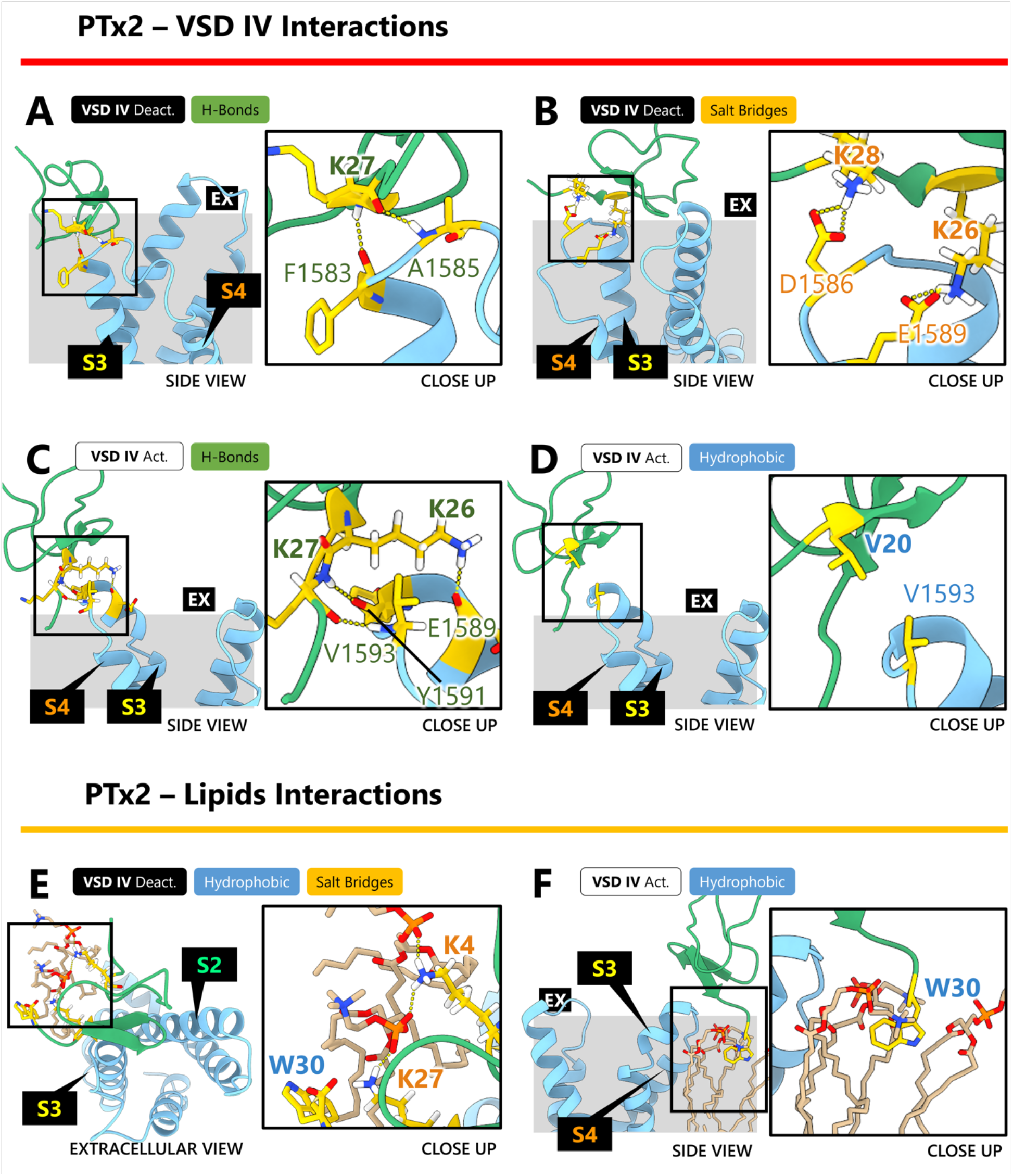
Dominant interactions recorded from molecular dynamics simulations between PTx2 and hNa_V_1.7 VSD IV (A, B, C, D) or PTx2 and POPC membrane (E, F). Residues engaged in binding are highlighted in gold. PTx2 is represented in green, while the VSD is depicted in blue. Each panel features interaction types presented in color-coded boxes whose colors correspond to the involved residues’ labels. The structures were visualized at the end of the 1-μs long MD simulations.

**Figure 7E** illustrates how the toxin bound to the deactivated VSD IV embedded itself within the membrane. PTx2 residues K4 and K27 generated salt bridges with lipid headgroups positioned proximal to the toxin’s binding site on the S3-S4 loop. At the same time, PTx2’s W30 initiated hydrophobic interactions with lipids, effectively functioning as a membrane anchor.

Beyond these dominant interactions, PTx2’s K26 was observed to form a hydrogen bond with VSD IV residue G1581, located lower in the S3 helix. Moreover, a limited number of interactions of PTx2 with the S1-S2 VSD IV residues were recorded. This includes hydrogen bonding between toxin and VSD IV residues T8 and E1534, hydrophobic interactions of L23 and W24 with Y1537 and V1541 respectively, and the formation of a salt bridge between R13 and E1534.

Upon binding with the activated VSD IV, as illustrated in **Figure 6D**, the toxin ascended significantly above the membrane, making only a limited number of interactions with residues on the S3-S4 loop. As represented in **Figure 6E**, the interactions established by the toxin with the activated VSD IV were predominantly hydrophobic (49 ± 3%) and hydrogen bonds (46 ± 2%), with salt bridges making up a minor portion (5 ± 1%). Throughout the simulations, the activated VSD IV underwent subtle conformational adjustments, as evidenced by a ∼2-3 Å RMSD relative to the initial structure, shown in **Figure 6F**. However, the binding site remained largely unchanged, exhibiting a minor ∼1.5 Å RMSD shift – a fluctuation nearly identical to that of the toxin itself.

With PTx2 situated atop the activated VSD IV S3-S4 loop, as depicted in **Figure 7C**, the toxin’s K26 side chain amino group formed a hydrogen bond with E1589 carbonyl group of the residue E1589 located on the S3 helix of VSD IV. Concurrently, toxin’s K27 backbone amide and carbonyl groups established dual hydrogen bonds with Y1591 carbonyl and V1593 amide groups, respectively, both positioned at the start of the VSD IV S3-S4 loop. Furthermore, PTx2’s V20 engaged in a hydrophobic interaction with VSD IV residue V1593, as visualized in **Figure 7D**. Despite PTx2’s significant elevation above the membrane, which resulted in substantial exposure to the solvent, its C-terminal tail reached down into the lipid bilayer. Here, toxin’s residue W30 engaged in hydrophobic interactions with nearby lipids in proximity to the VSD IV S3-S4 loop, maintaining sufficient toxin’s binding stability. Additionally, marginal hydrophobic contacts involving PTx2’s residue W5 and residue Y1591 on the VSD IV S3-S4 loop were detected.

### 2.3 Analysis of binding energetics

We calculated the free energy of binding between PTx2 and different states of hNa_V_1.7 VSD II and IV using the implicit-solvent MM/PBSA methodology^32^, based on our all-atom MD simulation trajectories. We analyzed each component of the free energy by block averages, taken during the latter half of each simulation (500 – 1000 ns) when binding stability was typically achieved as assessed by looking at the RMSD profiles (see **Figure 4** and **Figure 6**). These averages were calculated from all three replicas, as illustrated in **Figure 8**.

**Figure 8.**
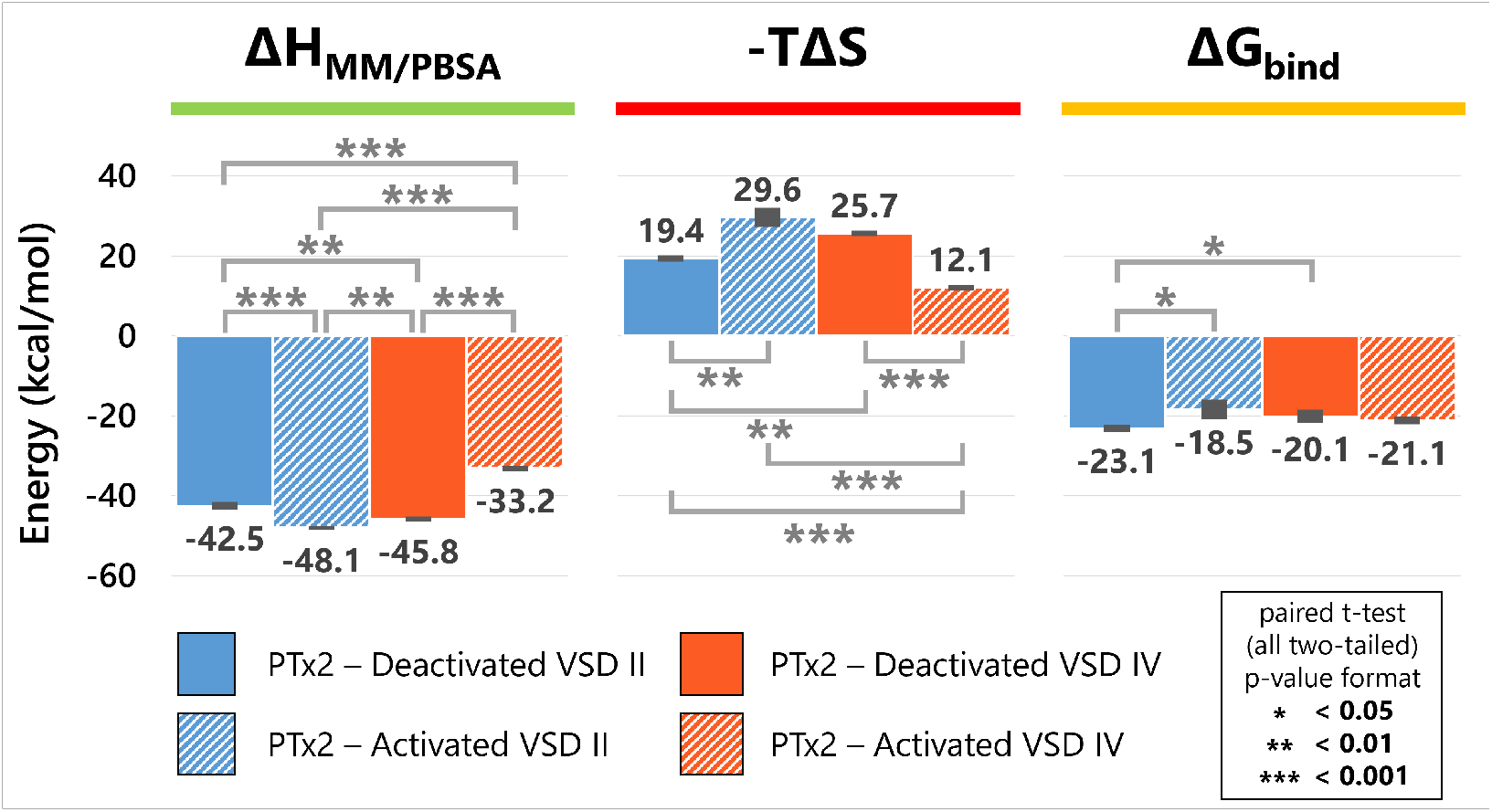
MM/PBSA binding energies in PTx2 – hNa_V_1.7 VSD systems averaged from three replicas from the second half (500 ns onward) of the all-atom molecular dynamics simulations. The optimal block sizes for the systems were selected to minimize the standard errors of the results.

The binding of PTx2 to the activated VSD II and deactivated VSD IV induced the most favorable binding enthalpy (ΔH_MM/PBSA_). In contrast, PTx2’s binding to activated VSD IV, marked by fewer intermolecular interactions in MD simulations, was characterized by the least negative enthalpy. When accounting for entropy (-TΔS), the resulting binding free energy of PTx2, ΔG_bind_, was most favorable for the deactivated VSD II, with a value of -23.1 ± 0.9 kcal/mol. This was followed by the activated VSD IV (-21.1 ± 1.0 kcal/mol), deactivated VSD IV (-20.1 ± 1.6 kcal/mol), and activated VSD II (-18.5 ± 2.5 kcal/mol) in descending order of favorability. Through paired t-tests, we established that a significant difference in Δ*G*_bind_ exists (*p < 0.05*) between PTx2 binding with the deactivated VSD II compared to the activated VSD II and the deactivated VSD IV.

Our calculations, showing that PTx2 binds more favorably to the deactivated state of VSD II, align with previous electrophysiology experiments^19,22^. Those studies demonstrated that PTx2’s antagonism of Na_V_1.7 occurs with around 60-fold reduced potency when the membrane potential is held at a voltage that favors VSD activation^22^. Moreover, another research study suggested that the estimated IC_50_ for PTx2’s inhibition of Na_V_1.7 activation (namely, binding to the deactivated VSD II) is roughly 400-fold lower than the observable IC_50_ for the inhibition of inactivation (that is, binding to the deactivated VSD IV)^19^.

Then we conducted a linear interaction energy analysis on our all-atom MD simulations using Amber20^33,34^. This allowed us to calculate the electrostatic and van der Waals interaction energies between PTx2 and VSD II/IV residues as well as explicit POPC lipids (not possible with the implicit-membrane MM/PBSA approach), and as depicted in **Figure 9** (for a more exhaustive depiction of specific interactions involving additional residues, refer to **Supplementary Figures S5-S7**).

**Figure 9.**
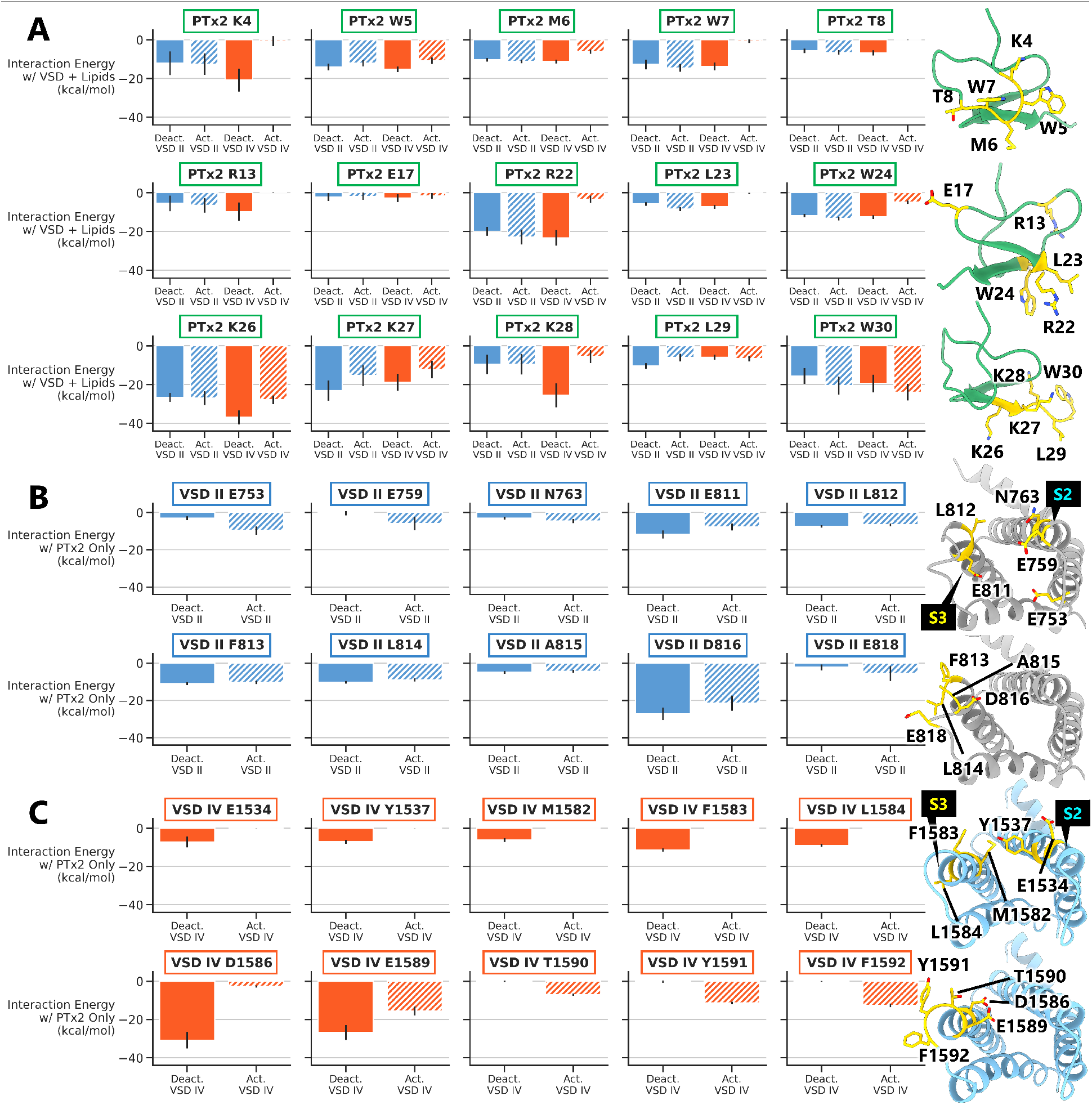
Total interaction energies contributed by each residue toward the binding process for different PTx2 – hNa_V_1.7 VSD MD systems. Residues involved in protein-protein interactions are visualized in yellow on the corresponding structure to the right of each row. The VSD structures are viewed from the extracellular side. The top 15 residues for PTx2 (A) and top 10 residues for VSD II (B) or IV (C) with the most favorable interaction energies (van der Waals + electrostatic) averaged from all three MD replicas are displayed.

**Figure 9A** shows the top 15 PTx2 residues that exhibit the most favorable total interaction energies with activated and deactivated VSD II/IV residues and lipids combined (to see the interaction energy with VSDs or lipids separately, refer to **Supplementary Figure S5**). The interaction energy for each toxin residue with different VSD states displays a variety of patterns. However, the general trend suggests that most residues have stronger interactions (i.e., more negative interaction energy values) with both the VSD II and VSD IV in their deactivated states compared to their activated states. This could imply that the electrostatic environment of the VSDs in their deactivated state is more favorable for binding these residues. From an energetic perspective, PTx2 residues R22, K26, K27, K28, and W30 are the main contributors to PTx2 – VSD II and IV binding through the formation of hydrogen bonds, hydrophobic interactions, and salt bridges. The most apparent difference is observed in the case of PTx2 binding to the activated state of the VSD IV as the binding pose excludes many toxin residues from forming interactions with the VSD.

Notably, toxin residues K4, R22, K26, K27, and K28 demonstrate particularly strong interaction energy with the VSDs. This could likely be due to their positive charge which might interact favorably with any negative residues of the VSDs and/or the POPC lipid PO4 groups. Conversely, PTx2’s E17 displays a much weaker interaction energy with the VSDs. Toxin residue K26 has notably stronger interactions with the deactivated VSD IV, showing the most favorable interaction energy observed in this data (-36.9 ± 3.6 kcal/mol), owing to its formation of a stable salt bridge with E1589.

The PTx2 residue K4 exhibits a significantly more favorable interaction energy when binding to the deactivated state of VSD IV compared to other cases. This can be attributed to the specific binding pose of the toxin in this state, which positions K4 favorably for forming salt bridges with surrounding lipids. The interaction energies with lipids depicted in **Supplementary Figure S5** revealed the main PTx2 – lipid membrane interaction surface, highlighting the important roles played by toxin residues K4, W5, M6, W7, W24, K27, and W30 in anchoring PTx2 to the cell membrane. The tight binding of PTx2 to the outer leaflet of the cell membrane seems to be critical for the inhibition of PTx2 on hNa_V_1.7 channels^26^. To support our findings, available mutagenesis studies that produced W5Y, W7Y, W24Y, and W30Y PTx2 mutant led to 6- to 291-fold reduction in potency in the inhibition of hNa_V_1.7^26^, indicating the importance of these residues for PTx2 binding to hNa_V_1.7 (in the cited study, the mutation to tyrosine was done since tyrosine is more polar than tryptophan and is less suited to bind at the water – lipid bilayer interface^35^).

Despite their significant roles in binding to both states of VSD II and the deactivated state of VSD IV, PTx2 residues K4, W7, T8, R13, R22, and L23 did not form substantial interactions when binding to the activated state of VSD IV. This lack of significant interactions could be attributed to the orientation of the bound toxin, which positioned these residues at a considerable distance from potential binding partners. As a result, their contribution to the binding process is diminished or negligible in this state.

**Figure 9B** provides an overview of the interaction energies between notable VSD II residues with PTx2. When comparing the activated and deactivated states, minor variations are observed in terms of the interaction energies of VSD II residues involved in the binding of PTx2. Notably, residues E811 and D816, located on the S3-S4 loop, exhibit slightly more favorable interaction energies with PTx2 when VSD II is in the deactivated state compared to the activated state. Conversely, residues E753 and E759 on the S1-S2 loop, as well as E818 on the S3-S4 loop, show stronger interactions with PTx2 when VSD II is activated compared to when it is in the deactivated state.

Interestingly, **Figure 9C**, which illustrates the interaction energies of notable VSD IV residues with PTx2, demonstrates a distinct difference in their pattern when PTx2 binds to the activated and deactivated states of VSD IV. This disparity in interaction energies can be attributed to the positioning of PTx2 within the membrane. In the deactivated state (**Figure 6A**), PTx2 binds deeper within VSD IV cleft, leading to a stronger interaction with the VSD residues. Conversely, in the activated state, PTx2 binds more superficially to VSD IV, resulting in fewer interactions and primarily involving VSD residues located at the top of the S3-S4 loop (**Figure 6D**). Additionally, the analysis of PTx2’s interactions with the lipid membrane reveals that when binding to VSD IV in the activated state, PTx2 exhibits significantly fewer interactions with the lipid membrane compared to the deactivated state (**Figure 7F** vs. **Figure 7B, D**). These findings highlight the nuanced differences in PTx2’s binding to VSD IV in the activated and deactivated states. The variations in interaction energies and the involvement of specific VSD residues, as well as the differential interactions with the lipid membrane, contribute to the distinct binding characteristics of PTx2 to VSD IV in different states.

## 3. Discussion

### 3.1 PTx2 interactions with hNa_V_1.7 VSD II and IV in different states

Using existing experimental data as a guide, we generated models of PTx2 bound to human Na_V_1.7 VSD II and IV in both the activated and deactivated states. These models accurately reproduced the crucial toxin-ion channel interactions at VSD II, as previously elucidated through mutational experiments and structural studies involving chimeric ion channels^17,22^. Our findings also shed new light on PTx2 interactions with VSD IV, unveiling substantial differences compared to its interactions with VSD II.

PTx2 primarily binds to VSD II in different states through interactions with residues on the S3-S4 loop of the VSD II. In both states, PTx2’s K27 formed hydrogen bonds with the F813 carbonyl and A815 amide located on the S3-S4 loop. In the deactivated state of VSD II, a hydrogen bond is established between residue K26 of PTx2 and residue E811 of VSD II. On the contrary, when VSD II is activated, K26 interacted with VSD II L812 instead, and the interaction was less frequent and more variable. Similarly, PTx2’s R22 established a more robust and consistent salt bridge with D816 (on the S3-S4 loop) when bound to the deactivated VSD II state compared to the activated state, as it occasionally shifted its interaction to E753 (on the S1-S2 loop). Also, PTx2 exhibited a hydrophobic interaction via L29 with L814 in the deactivated state of VSD II, an interaction not observed in the activated state.

PTx2 interacts with VSD IV primarily through residues on the VSD IV S3-S4 loop, but significant differences were observed in the toxin’s binding to the deactivated and activated states of VSD IV. For example, PTx2’s K26 formed a hydrogen bond with G1581 (located in the S3-S4 region) in the deactivated state and with E1589 (also in the S3-S4 region) in the activated state of VSD IV. In the activated state, K26’s interaction with E1589 was more frequent and consistent, suggesting a stronger bond. Moreover, the hydrophobic interaction between PTx2’s V20 and the VSD IV’s A1585, situated in the S3-S4 region, is weaker in the deactivated state than with V1593 in the activated state. Another differential interaction can be seen with the toxin’s residue K27. When binding to the activated VSD IV, K27 interacted with three different residues in the S3-S4 region, Y1591, V1593, and F1592, while in the deactivated state it interacted with F1583 and A1585 in S3-S4 region only.

### 3.2 PTx2 interactions with lipids when binding to hNa_V_1.7 VSD II and IV

A significant advancement highlighted in this study is the novel atomistic understanding of PTx2’s interactions with surrounding POPC membrane lipids upon binding to the Na_V_1.7 channel. Lipid interactions play a vital role in stabilizing the position and orientation of PTx2 for binding to VSD II and IV in different states. These novel insights from simulations could open a new avenue for enhancing PTx2 binding to the channel by optimizing toxin interactions with nearby membrane lipids^26^.

When VSD II is activated, PTx2 residue T8 formed a hydrogen bond with nearby POPC PO4 group, W30 engaged in hydrophobic interactions with the lipids’ fatty acid tails, and W7 formed cation-π interactions with POPC choline group. In contrast, when VSD II was deactivated, PTx2 residue W24 engaged in hydrophobic interactions while K27 formed salt bridges with POPC PO4 groups. The first view into the impacts of lipids to modulate toxin binding may constitute a novel and emerging area of research for optimizing peptide toxin’s specificity and selectivity.

When PTx2 binds to the activated state of VSD IV, the only interaction with lipids was a hydrophobic one originating from W30 which was stronger and more consistent than when PTx2 binds to the deactivated state. When PTx2 binds to the deactivated VSD IV, two salt bridges with POPC head groups were formed by toxin residues K4 and K27. It is noteworthy that K27 – POPC salt bridge interactions were substantially more frequent and consistent in the deactivated VSD IV than in the deactivated VSD II.

### 3.3 Energetics of PTx2 binding

Lastly, we complemented the data with specific binding energetic contributions from individual toxin and channel residues in different VSD states, enabling the identification of potential mutation candidates. This information can support the design of toxin analogs with enhanced potency, holding promise as a way to identify candidates for the development of innovative treatments for chronic pain. Our analysis of PTx2 binding energetics to different states of VSD II and IV indicates that PTx2 binds most favorably to the deactivated VSD II, followed by the activated VSD IV. A substantial difference in the free energy of PTx2 binding is observed when comparing the deactivated VSD II to both the activated VSD II and the deactivated VSD IV. PTx2 binding and trapping VSD II in its deactivated state can prevent the opening of the hNa_V_1.7 channel. Likewise, trapping VSD IV in its activated state can keep the channel in a non-conducting, inactivated state. The net effect is a decrease in the Na^+^ current flowing through the Na_V_1.7 channels when PTx2 is introduced, which aligns with experimental results^19^.

From a therapeutic standpoint, optimizing PTx2’s interactions with these states could enhance its effectiveness in pain therapy through hNa_V_1.7 channel inhibition. Furthermore, experimental findings demonstrate that the efficacy of PTx2 as an inhibitor of hNa_V_1.7 is significantly influenced by its binding to the lipid membrane^26^. Therefore, improving PTx2 interactions with cell membrane lipids could also augment PTx2’s inhibition of hNa_V_1.7.

Based on our analysis, we have identified certain residues of PTx2 that interact with both the VSDs (activated and deactivated states) and lipids but exhibit relatively weak interaction energies. As indicated in **Figure 9A** and **Supplementary Figure S5**, these residues include, but are not limited to, T8, D10, R13, E17, M19, V20, L23, W24, and L29. Specifically, T8, D10, R13, E17, and M19 primarily interacted with the lipid molecules in the membrane, with limited interactions observed only with the deactivated state of VSD IV. V20, on the other hand, interacted solely with the VSDs and did not form significant interactions with the lipids. However, residues L23, W24, and L29 demonstrated interactions with both the VSDs and the lipid molecules in the membrane.

By strategically mutating the identified residues, there is a possibility of enhancing the affinity of PTx2 towards the VSDs and/or lipids. These mutations can be designed to optimize specific interactions, such as salt bridges, hydrogen bonding, electrostatic, hydrophobic and/or other types of interactions, involved in PTx2’s binding to the VSDs and lipids as indicated in the contact maps in **Supplementary Figures S1-S4**. For example, considering that the toxin’s C-terminus tail primarily interfaces with lipid binding, introducing additional hydrophobic residues through mutations could enhance interaction with the fatty acid tails of the POPC lipids. Nonetheless, it is vital to balance this enhancement with the potential effects on state specificity and channel subtype selectivity for toxin-receptor binding, as well as the net effect on protein’s overall structure and stability.

### 3.4 Limitations and future directions

Through using computational modeling and simulation tools, we can model the atomic-resolution structures and relative energetics of peptide toxin interactions with hNa_V_1.7 in different conformational states. This allows for the accurate prediction of key binding interactions, their subsequent experimental validation and design of more potent and selective peptides targeting hNa_V_1.7 or other ion channels as we did recently^14^.

Although the computational workflow in our study can provide valuable insights into the complexities of peptide-receptor interactions at atomic resolution, several drawbacks are associated with each methodology. Given the absence of complete human channel structures with VSDs in the deactivated state, our modeling and simulations were limited to the VSDs of the channel. This limitation could potentially introduce instability during the simulations. Nonetheless, we observed only minor displacements of the PTx2 binding sites during the majority of our simulations as demonstrated by the RMSD profiles. In the future, we will expand our study by modeling and simulating full human channel structural models.

Peptide backbone flexibility, especially at the N- and C-termini, is difficult to capture during the protein-protein docking and, therefore, RosettaDock might not be able to sample the full range of possible toxin conformations. Additionally, Rosetta scoring uses an implicit membrane model that might not account for energetic contributions involving membrane lipids that could be important to stabilize the toxin pose. Although a lipid membrane can be explicitly included in MD simulations, simulation accuracy often depends on the quality of the force field and parameters used in the simulation, and it can be difficult to determine the best set of parameters for any given system^36^. Fully atomistic molecular dynamics simulations are limited by their timescale (ns to μs), and hence enhanced sampling MD simulation techniques need to be employed to observe large-scale protein conformational transitions^37,38^. For estimating the free energy of binding, MM/PBSA methodology may suffer from accuracy problems especially for predicting absolute binding free energies as was demonstrated by a study done by *Sheng et al.*^39^ The accuracy can be improved such as by damping the solvated and Coulombic terms for highly charged systems^39,40^, using residue type-specific dielectric constants^41^, and applying potentially more accurate entropy calculation methods^42^. Yet, MM/PBSA methodology employed here was successfully utilized in our recent study^43^ to predict conformation-dependent binding of a small molecule ligand norepinephrine and large stimulatory G protein to beta-2 adrenergic receptor, another integral membrane protein. Molecular simulation and analysis methodology advancements^38,44^ will be explored in follow-up studies with the current work providing a necessary framework and achieving a good agreement with several experimental findings^19,22^.

In conclusion, our molecular modeling and simulation protocol can be a helpful guide for designing more potent therapeutic peptide variants with improved selectivity for different states of the channel as well as other Na_V_ channel subtypes as will be explored in follow-up studies. In addition, our results demonstrate that the interactions between PTx2 residues and lipids are important for state-specific binding to Na_V_1.7 VSDs and should be considered in future channel inhibitor studies. These findings open the possibility to improve the balance of VSD and lipid-binding capability of the toxin and, consequently, its efficacy as a channel inhibitor. We anticipate that this methodology can be readily expanded to study the interactions between other peptides and ion channels. Advancing our understanding of the molecular mechanisms of ion channel modulation by peptide toxins will allow us to harness the full potential of these molecules for therapeutic applications.

## 4. Materials and methods

### 4.1 Structural modeling and peptide docking

In this study, we used computational modeling methods guided by experimental structural data to investigate molecular mechanisms of PTx2 interaction with the hNa_V_1.7 VSD II and VSD IV in multiple states (**Figure 10**). Our primary objective was to generate models of PTx2 bound to VSD II and VSD IV in deactivated and activated conformations as input for molecular dynamics (MD) simulations.

**Figure 10.**
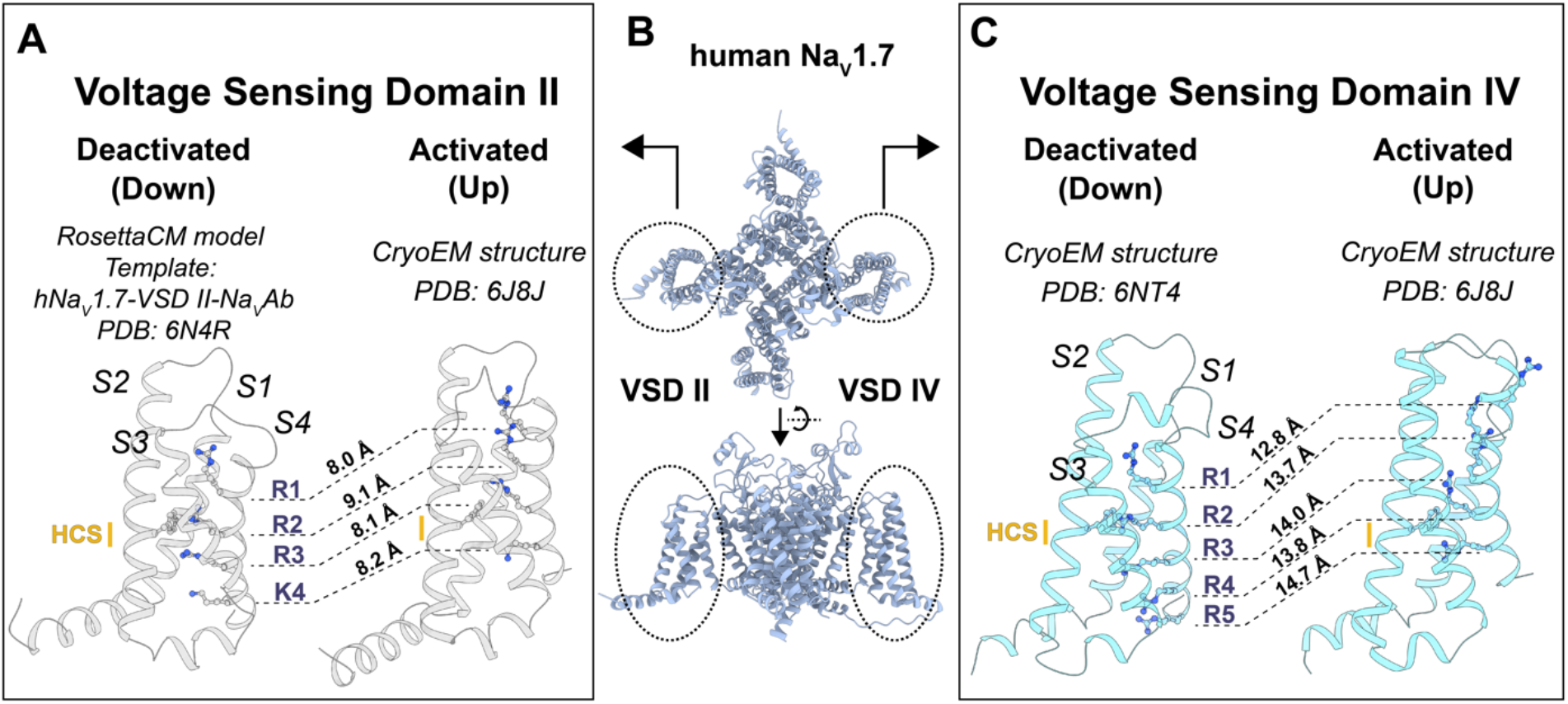
Structural comparison of different conformational states of hNa_V_1.7 VSD II and IV. A) Snapshots of deactivated (left) and activated (right) VSD II structures. For each state, S1 to S4 VSD segments are shown, highlighting the gating charge residues (“R1”, “R2”, “R3”, “K4”, “R5”) and the hydrophobic constriction site (HCS) in yellow. The distance between the C_α_ atoms of each gating charge in each state after superimposition of the VSDs is shown. B) Global view of cryo-EM Na_V_1.7 structure from PDB 6J8J^17^ highlights the VSD II and IV positions. C) Snapshots of deactivated (left) and activated (right) VSD IV structures (same information as in A).

To achieve this goal, we first used cryo-EM structures of PTx2 bound to hNa_V_1.7 to both the activated VSD II and VSD IV (PDBs: 6J8J^17^). However, the resolution of the toxin EM density, at ∼5 Å, did not permit complete atomic structure reconstruction in the experimental structure^17^. To overcome this limitation, we utilized the Rosetta structural modeling software suite^45^ and the PTx2 - hNa_V_1.7 complex electron-microscopy (EM) density data to model the interactions of PTx2 with activated hNa_V_1.7 VSD II and VSD IV. For VSD II, we used Rosetta Loop Modeling^46,47^ for modeling missing residues in the S3-S4 loop which form receptor site for binding of PTx2 based on structural^17^ and experimental data^22^.

The PTx2 docking protocol comprised a first step using RosettaDock^48,49^ with the Rosetta membrane energy function^50^. RosettaDock is a Monte Carlo-based multiscale algorithm that optimizes both rigid-body orientation and side chain conformation of the protein partners being docked. In this step, the ion channel VSD’s structural model was transformed into membrane coordinates by superimposing it into a reference hNa_V_1.7 experimental structure downloaded from the PDBTM database^51^ (which stores experimental structures of membrane proteins in membrane coordinates). The toxin was placed manually in three different initial locations around the binding site in the corresponding VSD, using the toxin’s EM density as a reference. The structure of PTx2 was obtained from the PTx2-hNa_V_1.7-VSD II-Na_V_Ab X-ray structure (PDB: 6N4I^22^). Subsequently, 20,000 docked models per input were generated with RosettaDock making a total of 60,000 models. The top 10% of models based on Rosetta *total_score* (total computed Rosetta energy of the protein) was extracted, and the top 100 *interface_score* (Rosetta interaction energy) models were used as an input for the next step. The second step involved EM density fitting of top-docked models to the experimental EM map using Rosetta EM density refinement^52^. During EM density refinement, toxin backbone and sidechain conformations are optimized using a modified energy function that accounts for the agreement between the model and the provided experimental EM map. In this step, 500 models were generated per input making a total of 50,000 models. The top 10% of models based on *total_score* was initially extracted. The final models were selected based on the recalculated *interface_score* after EM fitting, and on the *elec_dens_fast* score term (agreement between the Rosetta model and the experimental EM maps).

Furthermore, we docked PTx2 onto the structure of hNa_V_1.7 VSD IV trapped in a deactivated state by an α-scorpion toxin (PDB: 6NT4^4^), thereby constructing a model of PTx2 bound to deactivated hNa_V_1.7 VSD IV. Note that this experimental structure comprises a human - cockroach chimeric Na_V_ channel, in which the VSD IV has human sequence while the remaining part is derived from the cockroach Na_V_-PaS channel. To model the interactions between PTx2 and VSD IV, the Na_V_-PaS portion was removed, leaving only the VSD IV. The docking protocol was similar to the two cases above, but without any EM density-related refinement or scoring, since there is no experimental map available for this interaction. Finally, we employed the cryo-EM structure of the chimera channel hNa_V_1.7-VSD II-Na_V_Ab (PDB: 6N4R^22^), which has the VSD II in a deactivated state, as a template for RosettaCM^31^ homology modeling. This allowed us to generate a model of the deactivated VSD II with the entire hNa_V_1.7 VSD II sequence (sequence identity between template and target: 63.7%). We generated 5,000 RosettaCM models, extracted the top 10% based on *total_score*, and selected the final model based on both low RMSD for the template and low *total_score*. Then, we used RosettaDock^48,49^ with a membrane energy function^50^ to dock PTx2 to the deactivated VSD II of the hNa_V_1.7 model described above.

### 4.2 Atomistic MD simulations

Using the CHARMM-GUI web server^53^, the VSD-bound-toxin complexes were inserted into tetragonal phospholipid bilayer patches composed of 160 1-palmitoyl-2-oleoylphosphatidylcholine (POPC) molecules in total. The systems were then solvated by a 0.15 M aqueous NaCl solution to mimic physiological extracellular environment, resulting in molecular systems of ∼63,000 atoms as illustrated in **Figure 11**. The protonation state of each residue was assigned under neutral pH condition. Standard N- and C-termini were set for PTx2 and VSD II/IV. For PTx2, disulfide bonds were added between cystine residues C2-C16, C9-C21, and C15-C25.

**Figure 11.**
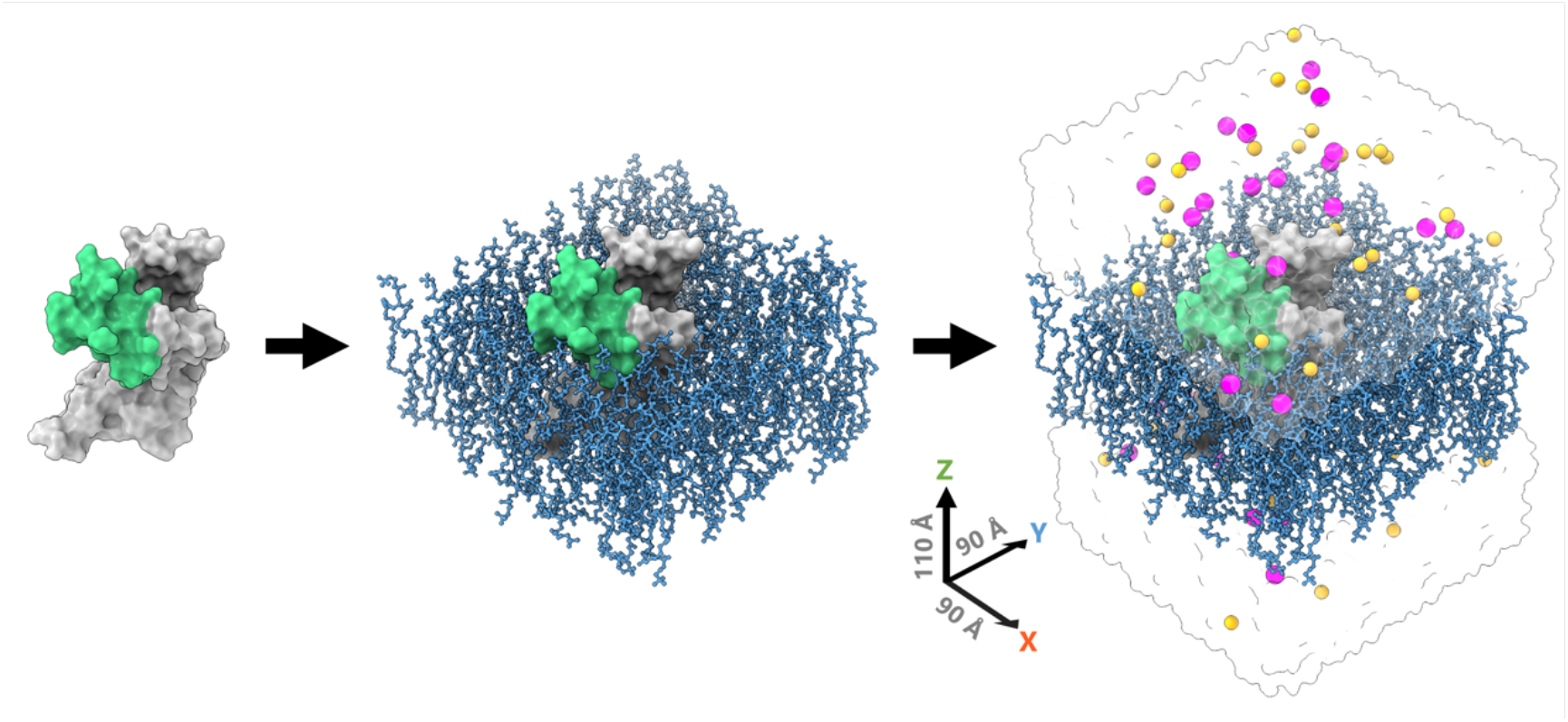
Creation process of a molecular system consisting of the PTx2 – hNa_V_1.7 activated VSD II complex embedded in a POPC bilayer and solvated with aqueous 0.15 M NaCl solution. PTx2 and VSD II are shown as green and dark gray surfaces, respectively. The lipid membrane composed of POPC lipids are shown in blue sticks. Water is displayed as a transparent surface. Na^+^ and Cl^-^ ions are displayed as purple and gold spheres, respectively.

All-atom molecular dynamics (MD) simulations were performed using the Amber20^33^ software package and the standard CHARMM all-atom force field for proteins (CHARMM36m), lipids (C36), ions^54–56^, and the TIP3P water model^57^. During system equilibration, a 1 kcal/mol/Å^2^ harmonic restraint was applied to all atoms in the system and was gradually reduced to 0.1 kcal/mol/Å^2^ over a period of 90 ns. Subsequently, a production run of 910 ns was conducted, resulting in a total simulation time of 1 μs. All MD simulations were run in the *NPT* ensemble at 310.15 K and 1 atm pressure maintained using the Langevin temperature equilibration scheme and Berendsen barostat. Non-bonded interactions were computed up to a 9 Å cutoff. Long-range electrostatic interactions were computed using Particle mesh Ewald (PME) method^58^, while no long-range correction was applied to van der Waals interactions as suggested for C36 force field. All covalent bonds involving hydrogen atoms were constrained using the SHAKE algorithm^59^ allowing to use a 2-fs time step. To maintain the structural stability of the VSD transmembrane helices (S1-S4), a weak 0.1 kcal/mol/Å^2^ harmonic restraint was imposed on their Cα atoms throughout the simulations. This restraint helped alleviate any structural instability arising from the isolation of the VSD from the full channel. The simulation protocol was replicated three times, each with different initial lipid positions and starting velocities of atoms. The replication accounts for potential variability in the lipid arrangement and system dynamics.

Afterward, protein-ligand interactions were characterized in each set of simulations using in-house Python scripts incorporating the protein–ligand interaction profiler (PLIP) Python module^60^. Binding free energy calculations were performed using the Molecular Mechanics Poisson-Boltzmann Surface Area (MM/PBSA) approach^61–63^ with all-atom simulation trajectories by MMPBSA.py program in Amber Tools^32^. MM/PBSA was selected due to its computational efficiency, which facilitates the generation of qualitative insights into the energetic components of binding. Moreover, it enables the incorporation of an implicit membrane in its calculations. This feature is particularly advantageous as it fosters a more accurate approximation of the energetics underlying the interaction between PTx2 and hNa_V_1.7, which is embedded in the membrane.

The Chamber module of ParmEd program was used to convert CHARMM-style forcefields to Amber-style forcefield^64^. The aqueous solution (with ionic strength of 0.15 M) and lipid membrane were treated implicitly using dielectric constants, denoted ε (water ε_w_ = 80, lipid bilayer ε_l_ = 2 and protein ε_p_ = 4). The solvent probe radius was set to 1.4 Å, and the atomic radii were set according to the converted force field parameters.

To obtain the solvation-free energy and the gas phase energy contributions without entropy, the particle-particle particle-mesh (P3M) procedure was used^65^. These calculations above were performed with implicit membrane, where the electrostatic energy includes both reaction field and Coulombic electrostatic energies. Entropy was calculated separately by the interaction entropy method^66^, where interaction energy between PTx2 and hNa_V_1.7 VSD II/IV including only Coulombic electrostatic and van der Waals energies was needed. To obtain the Coulombic energy separated from the reaction field energy, each system energy was recalculated using the dielectric boundary surface charges method in the implicit ionic solution. To calculate the energy contribution of each residue to the binding process for each system, we computed the electrostatic and van der Waals interaction energies of PTx2 with POPC lipids and VSD II/IV using AMBER linear interaction energy analysis^33,34^.

### 4.3 Molecular graphics

Molecular graphics visualization was performed using UCSF ChimeraX^67^ to depict all resulting models and illustrate toxin – VSD protein residue interactions. Structural comparison was carried out through the superposition of the structures using the MatchMaker tool within ChimeraX.

## 5. Supplementary Information

Supplementary information (SI) includes: 1) PTx2, hNa_V_1.7 VSD II and VSD IV amino acid sequences used in this study; 2) SI Figures S1-S7 detailing specific PTx2 - hNa_V_1.7 VSD/lipids inter-residue interactions including their frequencies and interaction energies; 3) two SI videos demonstrating structural transitions of PTx2 - hNa_V_1.7 VSD II and VSD IV complexes between deactivated and activated VSD states using Rosetta Dock models.

## Acknowledgments

We would like to thank all members of the I.V., C.E.C. and V.Y.-Y. laboratories for helpful discussions. This work was supported by National Institutes of Health Common Fund Grant OT2OD026580 (to C.E.C. and I.V.), National Heart, Lung, and Blood Institute (NHLBI) grants R01HL128537, R01HL152681, and U01HL126273 (to C.E.C., V.Y.-Y. and I.V.), American Heart Association Career Development Award grant 19CDA34770101 (to I.V.), National Science Foundation travel grant 2032486 (to I.V.), UC Davis Department of Physiology and Membrane Biology Research Partnership Fund (to C.E.C. and I.V.) as well as UC Davis T32 Predoctoral Training in Basic and Translational Cardiovascular Medicine fellowship supported in part by NHLBI Institutional Training Grant T32HL086350 (to K.N.) and UC Davis Chemical Biology Program fellowship supported in part by National Institute of General Medical Sciences (NIGMS) Institutional Training Grant 5T32GM136597-02 (to K.C.R.). Computer allocations were provided through Extreme Science and Engineering Discovery Environment (XSEDE) grant MCB170095 (to I.V., C.E.C., and V.Y.-Y.), Texas Advanced Computing Center (TACC) Leadership Resource and Pathways Allocations MCB20010 (to I.V., C.E.C., and V.Y.-Y.), Oracle cloud credits and Oracle for research fellowship award (to I.V., C.E.C. V.Y.-Y.), Pittsburgh Supercomputing Center (PSC) Anton 2 allocations PSCA17085P, MCB160089P, PSCA18077P, PSCA17085P, PSCA16108P (to I.V., C.E.C., and V.Y.-Y.). Anton 2 computer time was provided by the Pittsburgh Supercomputing Center (PSC) through Grant R01GM116961 from the National Institutes of Health. The Anton 2 machine at PSC was generously made available by D.E. Shaw Research.

