## Supplemental text, tables and figures for "Elucidating Molecular Mechanisms of Protoxin-2 State-specific Binding to the Human Na_V_1.7 Channel"

for

**Running title:** Protoxin-2 - $Na_V$1.7 channel state-specific binding

### Supplementary Text

*Amino acid sequences of the structures used in the study.*

*Multiple sequence alignment was done using CLUSTAL OMEGA (1.2.4)*

```
Protoxin2                         YCQKWMWTCDSERKCCEGMVCRLWCKKKLW


hNaV1.7_VSD_II_deactivated        CSPYWIKFKKCIYFIVMDPFVDLAITICIVLNTLFMAMEHHPMTEEFKNVLAIGNLVFTG        60
hNaV1.7_VSD_II_activated          CSPYWIKFKKCIYFIVMDPFVDLAITICIVLNTLFMAMEHHPMTEEFKNVLAIGNLVFTG        60
hNaV1.7_VSD_IV_deactivated        ------KIQGCIFDLVTNQAFDISIMVLICLNMVTMMVEKEGQSQHMTEVLYWINVVFII        54
hNaV1.7_VSD_IV_activated          ----GNKIQGCIFDLVTNQAFDISIMVLICLNMVTMMVEKEGQSQHMTEVLYWINVVFII        56
                                          *:: **: :* :  .*::* : * ** : * :*:.  ::.:.:**   *:**

hNaV1.7_VSD_II_deactivated        IFAAEMVLKLIAMDPYEYFQVGWNIFDSLIVTLSLVELFLADVEGLSVLR--SFRLLRVF        118
hNaV1.7_VSD_II_activated          IFAAEMVLKLIAMDPYEYFQVGWNIFDSLIVTLSLVELFLADVEGLSVLR--SFRLLRVF        118
hNaV1.7_VSD_IV_deactivated        LFTGECVLKLISLRHY-YFTVGWNIFDFVVVIISIVGMFLADLIETYFVSPTLFRVIRLA        113
hNaV1.7_VSD_IV_activated          LFTGECVLKLISLRHY-YFTVGWNIFDFVVVIISIVGMFLADLIETYFVSPTLFRVIRLA        115
                                    :*:.* *****::  * ** ******* ::* :*:* :****:    .:    **::*

hNaV1.7_VSD_II_deactivated        KLAKSW------  124
hNaV1.7_VSD_II_activated          KLAKSW------  124
hNaV1.7_VSD_IV_deactivated        RIGRILRLV---  122
hNaV1.7_VSD_IV_activated          RIGRILRLVKGA  127
                                  :::.:
```

### Table S1. *Criteria Used for Detection of Non-bonded Interactions.*

| TYPES OF INTERACTION | DETECTION CRITERIA | FILTERING CRITERIA |
|---|---|---|
| **Hydrophobic Interaction** | All pairs of non-bonded atoms not involved in other specific types of interactions shown below within a distance of 4 Å. | Exclude hydrophobic interactions between rings that interact through the π-stacking. |
| **Hydrogen Bonds** | Distance between the hydrogen bond donor (D) and acceptor (A) should be less than 3.5 Å.<br>The angle D-H···A should be above 100°. | Exclude hydrogen bonds between atoms that already form a salt bridge.<br>A hydrogen bond donor can participate in only one hydrogen bond, while acceptor atoms can form multiple hydrogen bonds. |
| **Salt Bridges** | Centers of oppositely charged groups that come within a distance of 5.5 Å. | |
| **Cation-π** | Pairing of a positive charge and an aromatic ring if the distance between the charge center and the aromatic ring center is less than 6 Å. | |
| **π-Stacking** | Centers of two aromatic rings within a distance of 5.5 Å. The angle between the rings should deviate no more than 30° from the optimal angle (90° for T-stacking, 180° for P-stacking). When projecting each ring center onto the opposite ring plane, the distance between the other ring center and the projected point (offset) should be less than 2 Å. | |

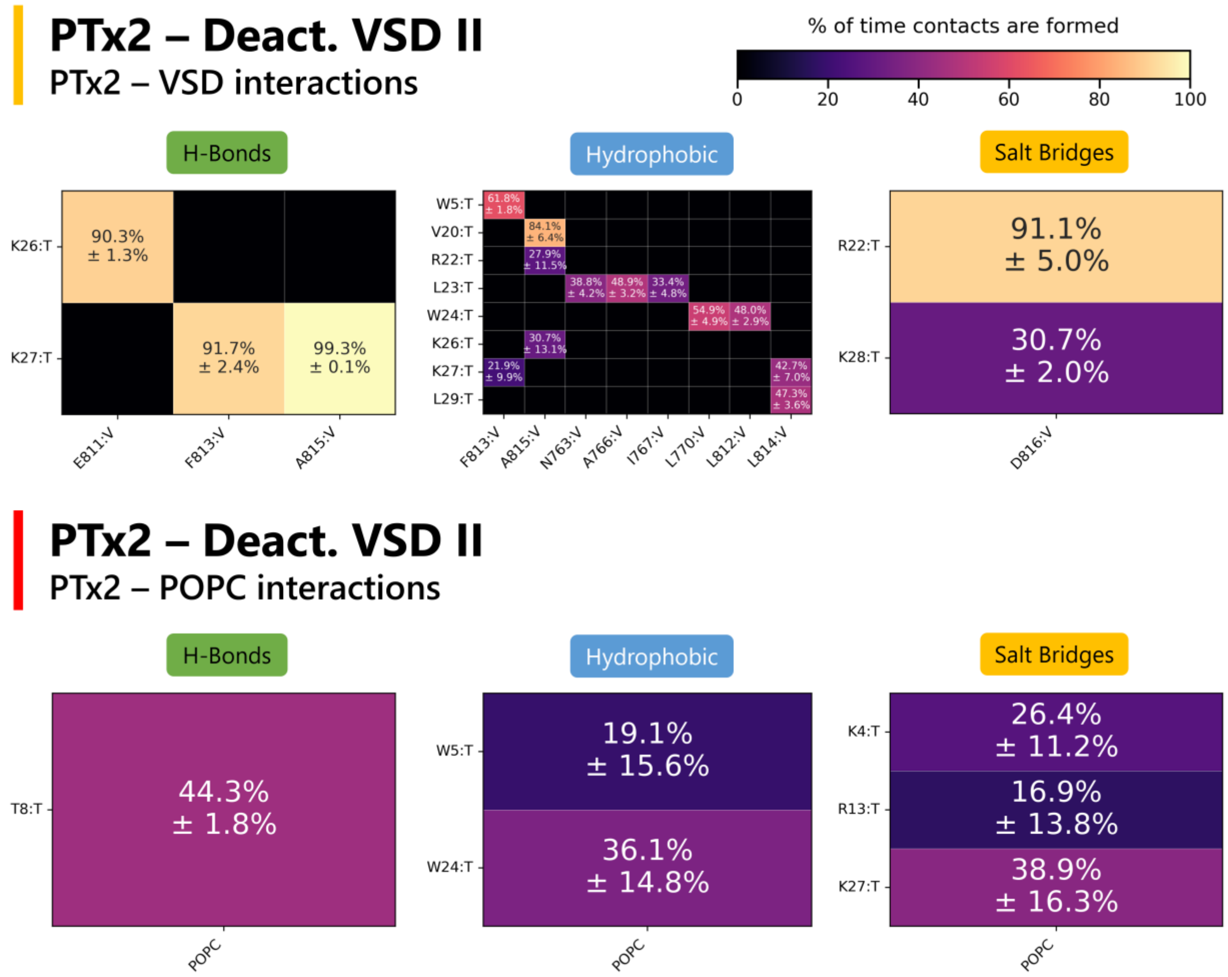


**Figure S1.** ***Contact maps showcasing PTx2 – VSD residues or POPC lipid interactions averaged from three MD simulation replicas of PTx2 – deactivated VSD II.*** *PTx2 residues are denoted by the suffix ":T", while VSD residues are denoted by the suffix ":V" Error bars are standard errors of mean from 3 MD replicas.*

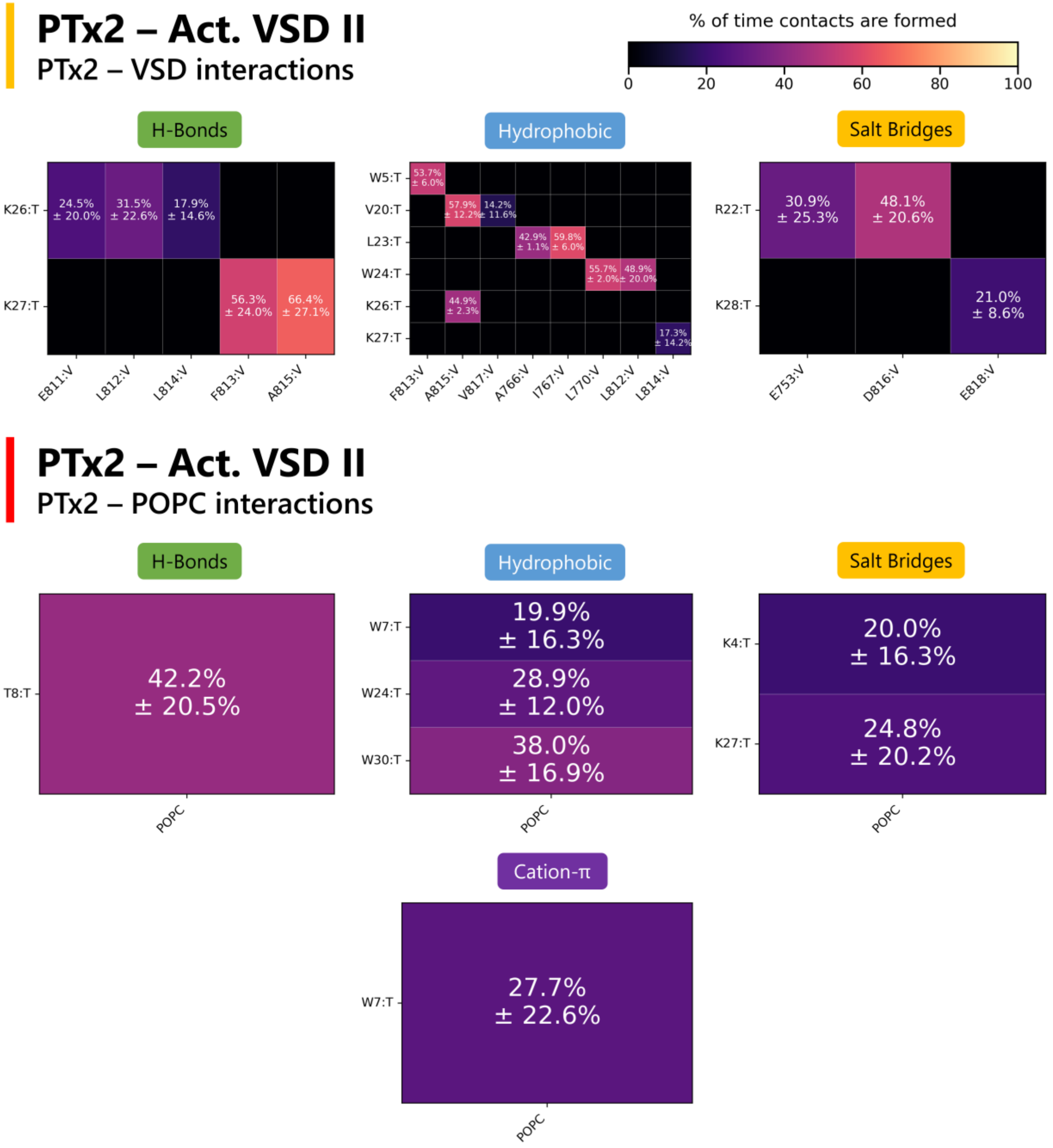


**Figure S2.** ***Contact maps showcasing PTx2 – VSD residues or POPC lipids interactions averaged from three MD simulation replicas of PTx2 – activated VSD II.*** *PTx2 residues are denoted by the suffix ":T", while VSD residues are denoted by the suffix ":V". Error bars are standard errors of mean from 3 MD replicas.*

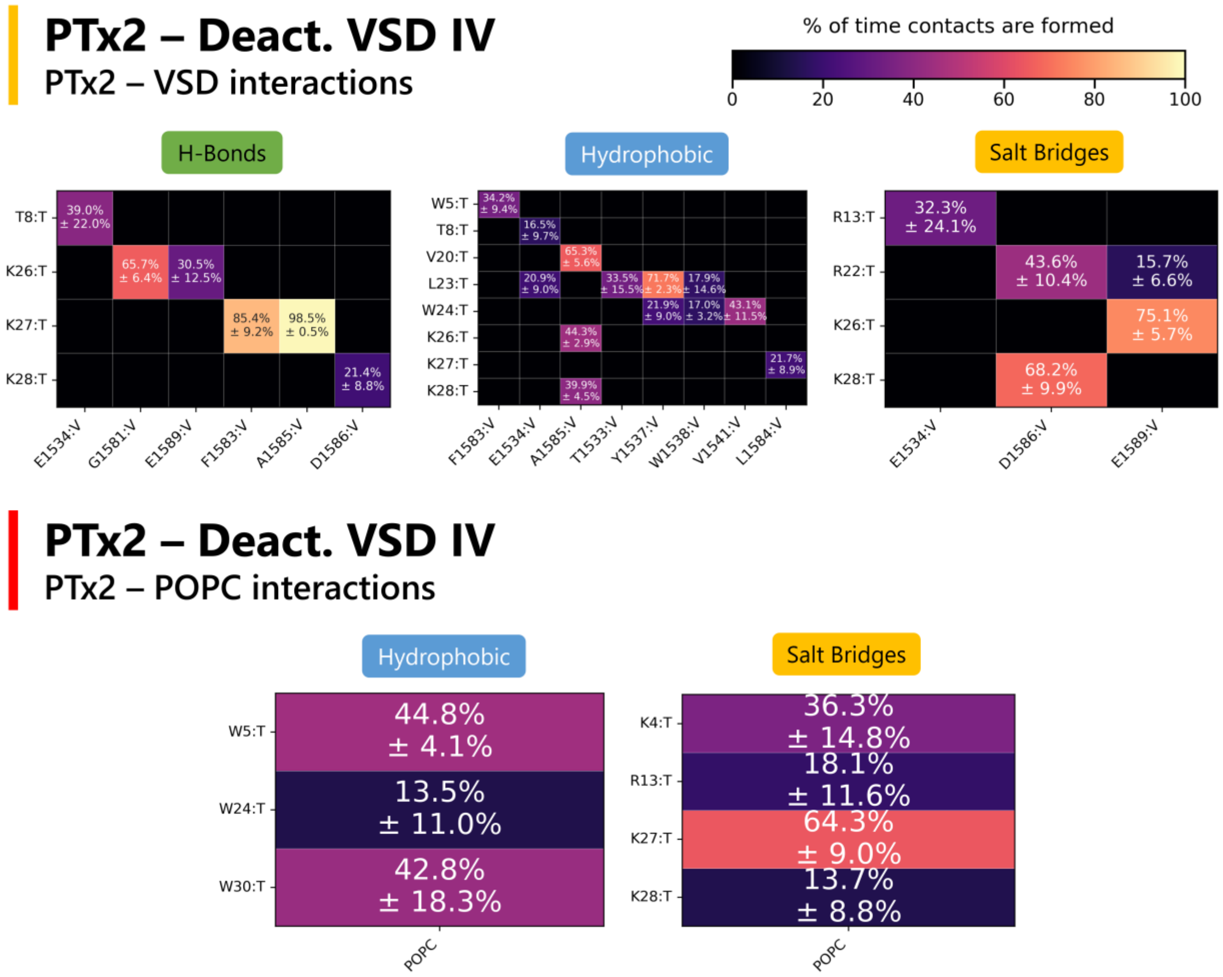


**Figure S3.** ***Contact maps showcasing PTx2 – VSD residues or POPC lipids interactions averaged from three MD simulationreplicas of PTx2 – deactivated VSD IV.*** *PTx2 residues are denoted by the suffix ":T", while VSD residues are denoted by the suffix ":V". Error bars are standard errors of mean from 3 MD replicas.*

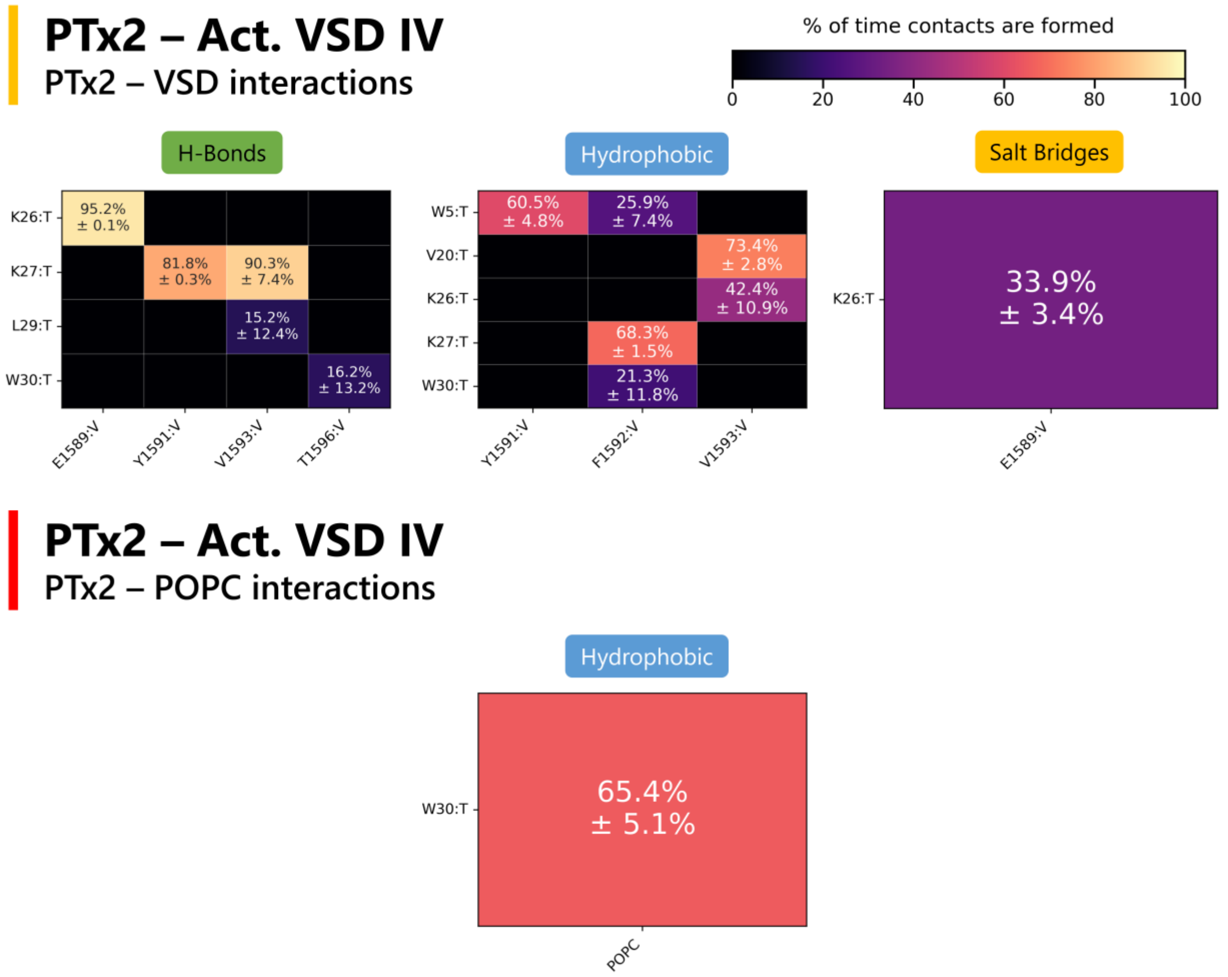


**Figure S4.** ***Contact maps showcasing PTx2 – VSD residues or POPC lipids interactions averaged from three MD simulation replicas of PTx2 – activated VSD IV.*** *PTx2 residues are denoted by the suffix ":T", while VSD residues are denoted by the suffix ":V". Error bars are standard errors of mean from 3 MD replicas.*

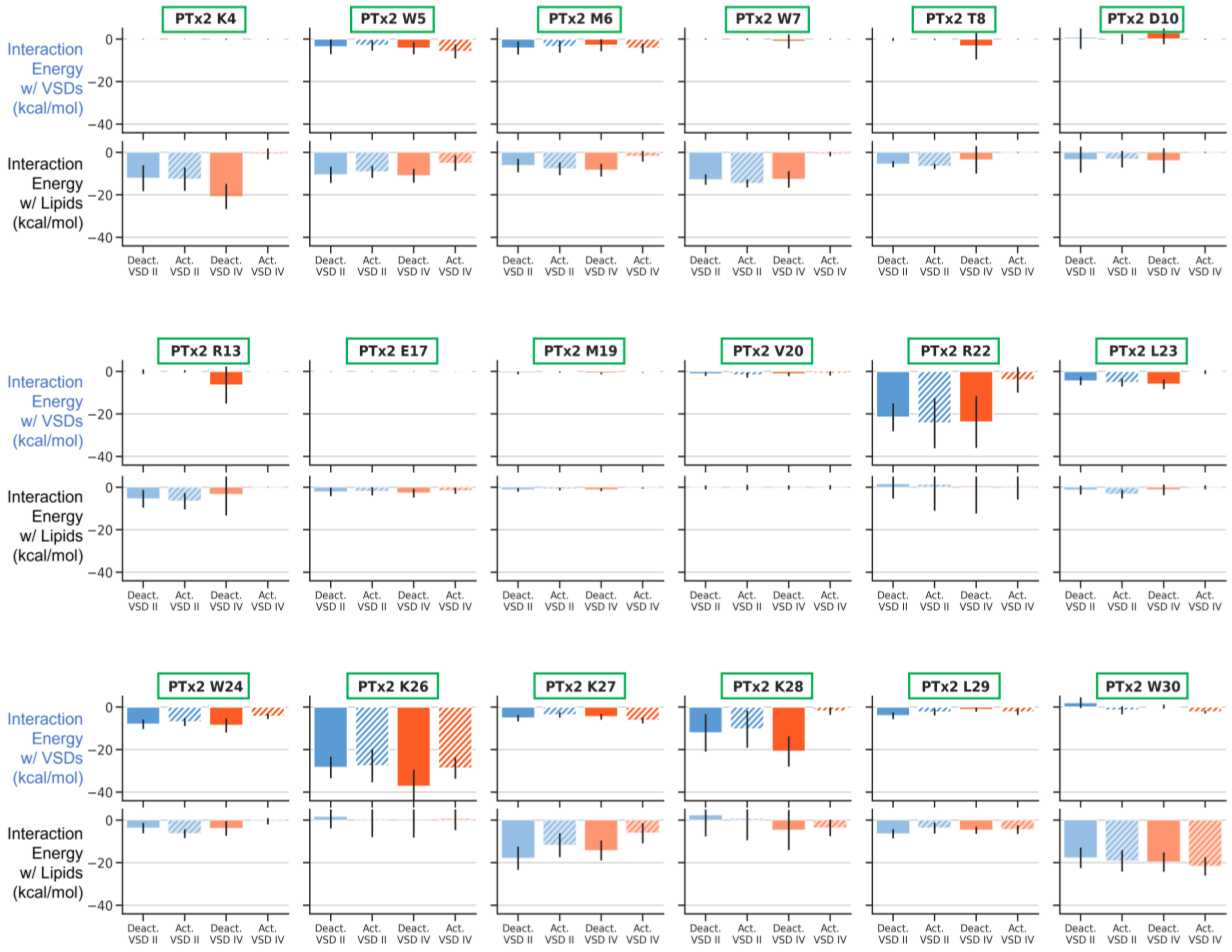


**Figure S5.** ***Interaction energies (van der Waals + electrostatic) contributed by each PTx2 residue toward the VSD and lipid binding process for different PTx2 - hNa$_V$1.7 VSD II/IV*** **MD simulation** ***systems.*** *Only residues with combined interaction energies across all systems more favorable than -5 kcal/mol when averaged from all replicas are displayed.*

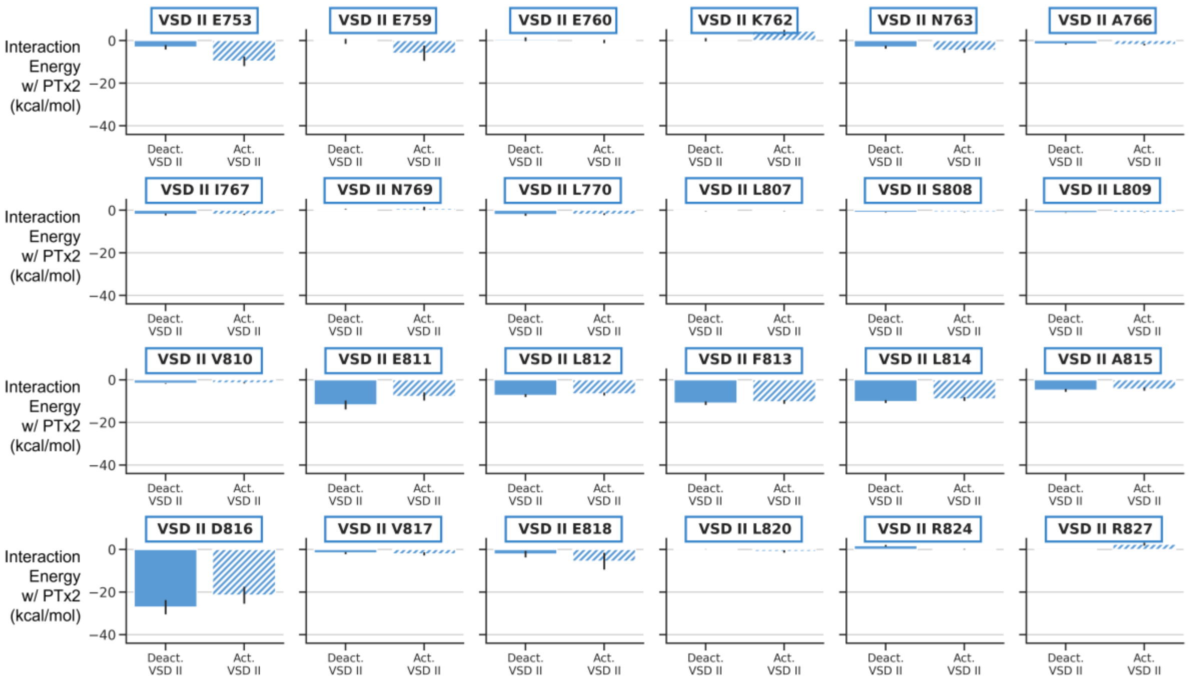


**Figure S6.** ***Interaction energies (van der Waals + electrostatic) contributed by each VSD II residue toward the PTx2 binding process for different PTx2 - hNa$_V$1.7 VSD II/IV MD simulation systems.*** *Only residues with combined interaction energies across all systems more favorable than -0.5 kcal/mol when averaged from all replicas are displayed.*

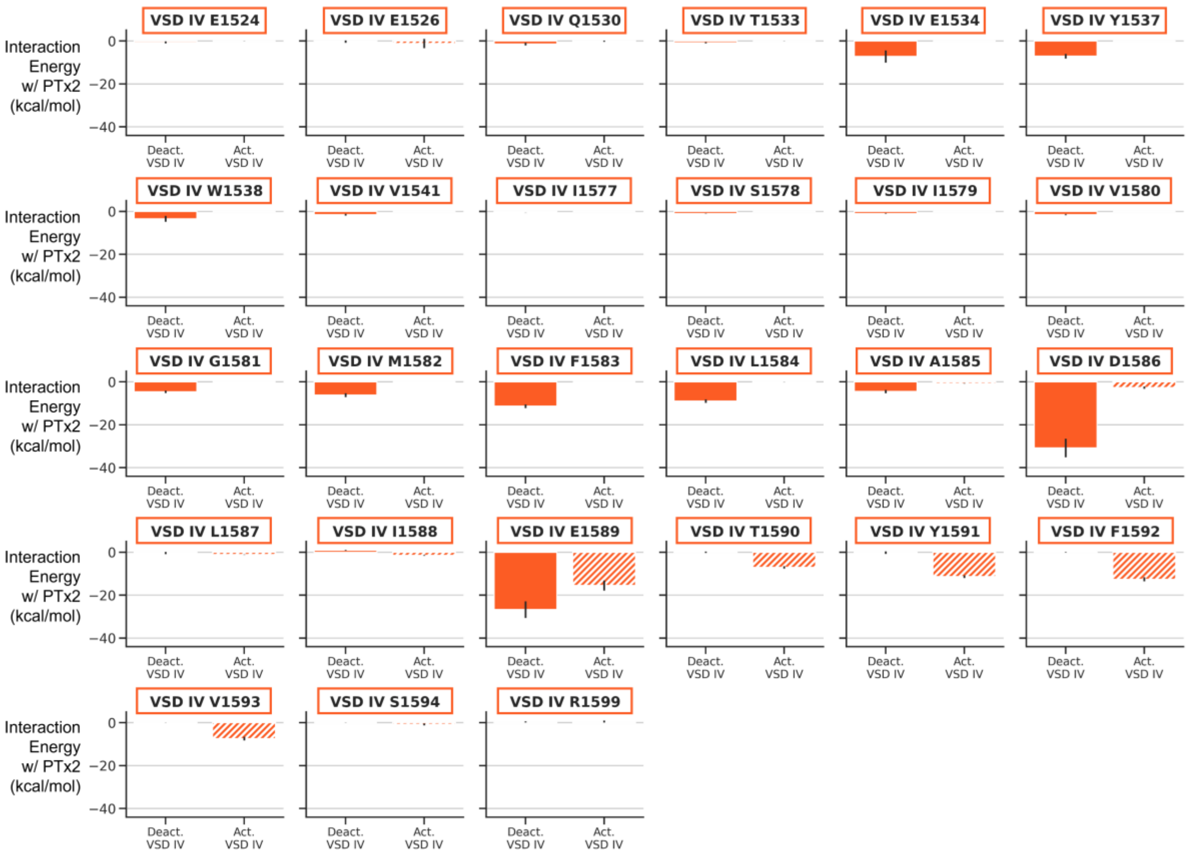


**Figure S7.** ***Interaction energies (van der Waals + electrostatic) contributed by each VSD IV residue toward the PTx2 binding process for different PTx2 - hNa$_V$1.7 VSD II/IV MD simulation systems.*** *Only residues with combined interaction energies across all systems more favorable than -0.5 kcal/mol when averaged from all replicas are displayed.*
